# Signaling mechanism of the transmembrane energy receptor Aer

**DOI:** 10.64898/2026.05.25.727702

**Authors:** Flory A. Olsthoorn, Alise R. Muok, Zachary A. Maschmann, Yajie Xu, Siddarth Chandrasekaran, Robert Dunleavy, Brian R. Crane

**Affiliations:** Department of Chemistry and Chemical Biology, Weill Institute for Cell and Molecular Biology, Cornell University, Ithaca, NY 14853; Structural Biochemistry, Bijvoet Center for Biomolecular Research, Utrecht University, Utrecht, The Netherlands, 3584 CG; Department of Molecular Biology, Princeton University, Princeton, NJ 08544; Intrepid Labs, Toronto, Canada

**Author notes:** Correspondence: Brian R. Crane.

**Keywords:** Chemotaxis, Energy Taxis, Redox Sensing, Transmembrane Signaling, Cryo-EM DEER Spectroscopy, Flavoprotein

## Abstract

The *E. coli* aerotaxis receptor Aer is a bacterial chemoreceptor that senses intracellular redox changes via an N-terminal PAS domain bound to a flavin adenine dinucleotide (FAD) cofactor. Distinct from canonical methyl-accepting chemotaxis proteins (MCPs) such as Tar/Tsr, Aer lacks a periplasmic ligand-binding domain and adaptive methylation, transmitting conformational signals laterally from the PAS domain to the HAMP domain and the methylation helix-like cap (MHL-cap) of the kinase control domain (KCD). To elucidate the Aer signalling mechanism, we determined cryo-electron microscopy (cryo-EM) structures of full-length Aer in oxidized flavin quinone (kinase-on) and reduced semiquinone (kinase-off) states. Structural comparison reveals redox-linked rearrangements of the FAD-binding pocket, reorientation of PAS–HAMP interactions, and strikingly altered MHL-cap stability. PAS–MHL-cap contact in the oxidized state compresses the receptor and stabilized proximal KCD helices, whereas reduction disrupts these contacts, increasing KCD flexibility. To probe distal effects on KCD architecture, we performed nanodisc reconstitution and pulse dipolar ESR spectroscopy on spin-labelled positions along the four-helix bundle. Distance distributions indicate redox-dependent changes in helix separation, particularly at the C-terminal MHL2 region, consistent with PAS-driven loosening of KCD packing in kinase-off states. These data support a model in which FAD redox chemistry reorganizes flavin pocket residues that in turn subtly alter PAS conformation to influence PAS-HAMP and PAS-MHL-cap packing and hence KCD conformational stability. The findings reveal an Aer-specific signaling axis distinct from periplasmic-ligand binding MCPs that has adapted MCP architecture for lateral PAS input and cytoplasmic redox sensing.

**Significance:** Transmembrane energy receptors, such as *E. coli* Aer, interpret metabolic signals to optimally position bacteria in their environment. We apply cryo-EM and pulse-diplolar ESR spectroscopy to characterize the structural landscape of full-length Aer in oxidized and reduced states to reveal a signal transmission mechanism distinct from that of canonical methyl-accepting chemotaxis proteins (MCPs). This work establishes Aer as a specialized MCP variant adapted for cytoplasmic redox sensing through a flavin-containing PAS domain that yet displays features of conformational propagation long suspected to be critical for signaling by the greater family of MCP transmembrane receptors. These findings open possibilities for engineering redox- and light-responsive receptors and targeting energy taxis pathways in pathogens.

## Introduction

Bacteria navigate their environment through a receptor-mediated behavior known as chemotaxis^1^. Through the sensing of chemical stimuli, bacteria localize to beneficial microenvironments for optimal metabolism, growth and pathogenicity. Closely related to chemotaxis, energy taxis involves motility responses to indirect changes in metabolic state, mediated by the availability of high-energy nutrients, cellular redox status, and respiration^2–4^. For example, a class of transmembrane receptor typified by *E. coli* Aer (Fig. 1) detect changes in terminal electron acceptors of the electron transport chain (ETC) and couple outputs to the chemotactic signaling infrastructure to control flagellar rotation bias^4–6^. Certain reactive oxygen species, such as peroxide, also drive Aer-mediated repellent responses, which are linked to antibiotic avoidance^7^.

**Figure 1:**
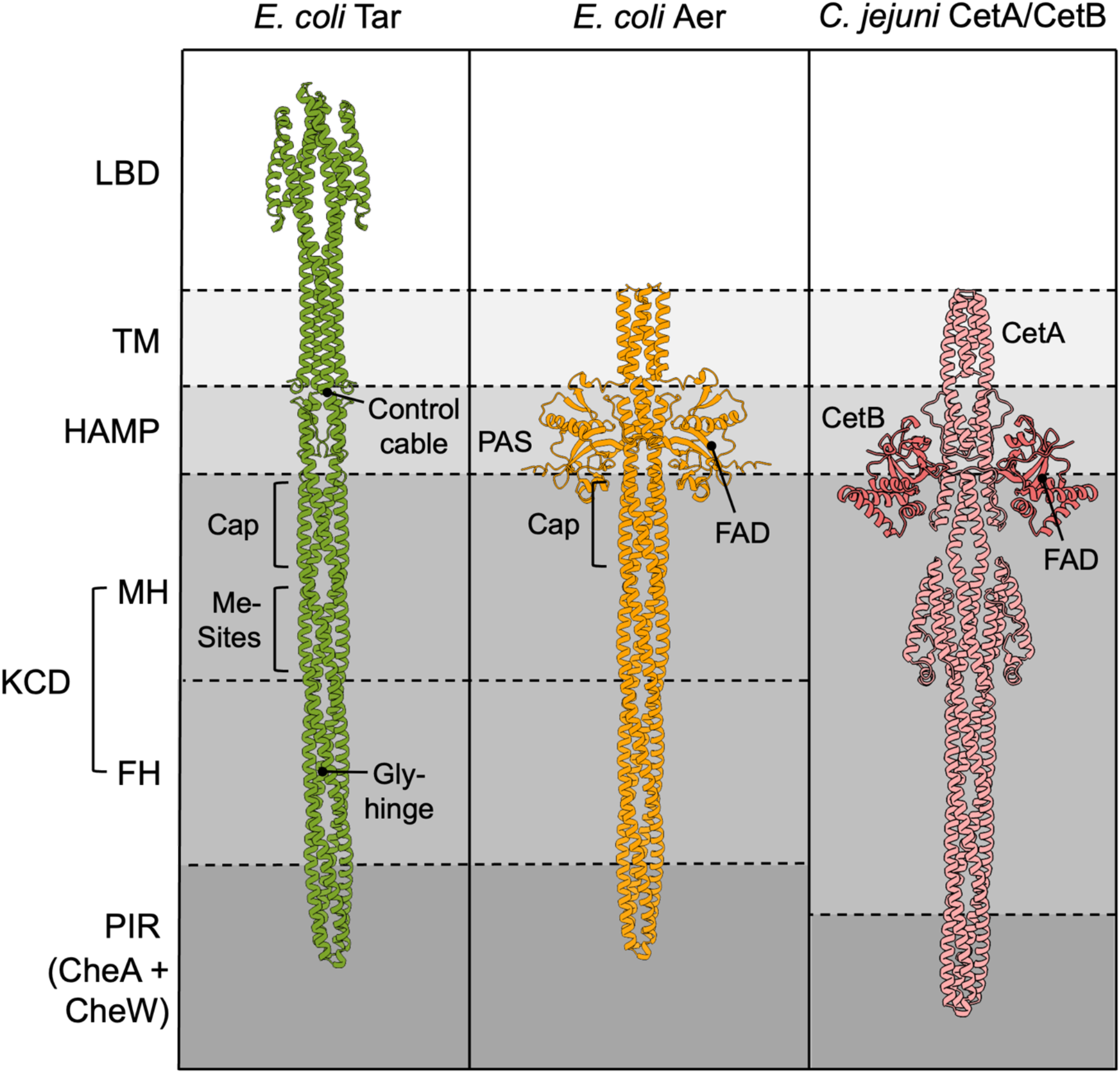
Three classes of bacterial receptors. AlphaFold 3 predictions of bacterial receptors *E. coli* Tar, *E. coli* Aer, and *C. jejuni* CetA/CetB illustrating different architectures of receptor signalling. *E. coli* Tar represents a canonical MCP with a periplasmic ligand binding domain (LBD), transmembrane region (TM), Histidine kinase, Adenylate cyclase, Methyl-accepting chemotaxis protein and Phosphatase (HAMP) domain, kinase control domain (KCD) made up of the methylation helix cap (MH-cap) region, which contains conserved Glu residues that undergo reversible methylation by CheR and CheB (Me-Sites), the flexible helix region (FH) that contains the glycine hinge (Gly hinge) and the protein interaction region (PIR) that binds CheA and CheW. *E. coli* Aer has a redox sensing PAS domain instead of an LBD and does not undergo adaptational methylation. *C. jejuni* CetA/CetB structurally resembles Aer, except its PAS module (CetB, pink) is a separate protein from the signalling unit (CetA, salmon).

Chemoreceptors, including Aer, are dimeric transmembrane proteins of the methyl-accepting chemotaxis protein (MCP) class, named for their ability to fine-tune sensory output through reversible methylation of specific cytoplasmic residues (Fig. 1)^8, 9^. In most types of MCPs, sensing occurs through ligand binding events at the periplasmic ligand binding domain (LBD) that induce local conformational changes that are transmitted through the membrane to the cytoplasmic receptor tip^10^. Cytoplasmic signal propagation is mediated by highly conserved coiled-coil modules: the HAMP (Histidine kinases, Adenyl cyclases, Methyl-accepting proteins, Phosphatases) domain, which forms a parallel 4-helix bundle^11^, and the kinase-control module (KCD), which forms an anti-parallel 4-helix bundle. A protein interaction region (PIR) at the KCD tip interacts with the histidine kinase CheA and coupling protein CheW to form receptor:kinase arrays of trimers-of-receptor dimers and hexagonal symmetry near the cell poles. The PIR regulates CheA autophosphorylation activity in a ligand-dependent manner^12, 13^. CheA transfers phosphate to activate CheY, which then directly interacts with the flagellar motor and influences its rotation sense^1^. Aer-type receptors are also obligate homodimers but differ from conventional MCPs in structure and function^6, 14, 15^. Aer contains a transmembrane region (TM), HAMP and C-terminal KCD, but instead of a periplasmic LBD, the sensing module is an N-terminal PAS (Per-Arnt-Sim) domain^16, 17^ that couples to the TM via an F1 linker^18^ (Fig. 2A). The F1 linker connecting the PAS domain to the TM region is not directly involved in signalling, but supports the maturation of the PAS and HAMP modules^18–20^. In some energy receptors, such as *C. jejuni* CetA/B, the PAS module is a separate protein^21^ (Fig. 1).

**Figure 2:**
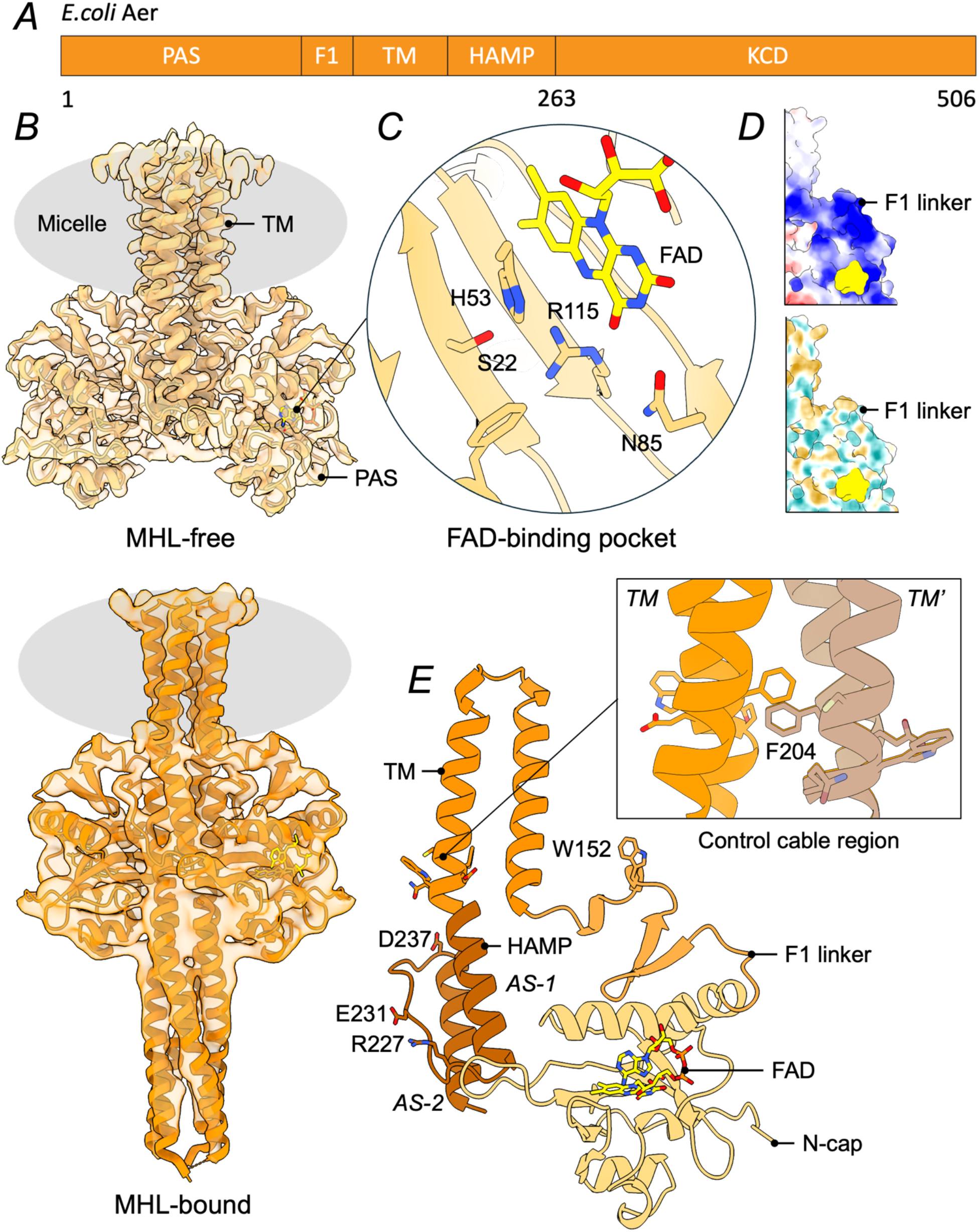
Oxidized Aer adopts two conformations in solution: MH-free and MH-bound. (A) Sequence schematic of *E. coli* Aer, designating the positions of the PAS domain, F1 linker, transmembrane helices, HAMP helices, and KCD helices. (B) Cartoon of the MH-free (top, tan) and MH-bound (bottom, orange) conformations of Q Aer fit into their respective electron density maps (LMNG micelle area, gray). (C) Details of the FAD-binding pocket of MH-free showing key residues that interact with the FAD isoalloxazine ring (yellow). (D) Surface representation of the F1-linker region of the Aer MH-bound conformation colored according to the electrostatic potential (blue -positive potential, red - negative) or hydrophobicity (tan - hydrophobic, cyan – hydrophilic). FAD is shown in yellow. (E) Cartoon of MH-free monomeric subunit colored by PAS, F1-linker, TM, and HAMP domains. W152 on F1 inserts into the micelle, and E231-R227 form a salt bridge on the HAMP connector. D237 of the DExG motif caps the N-terminus of the AS-2 helix. Inset shows the residues of the control cable in context of the dimeric TM module.

The conformational changes of chemoreceptors that propagate signals between the periplasm and the cytoplasm have garnered much attention^10, 22, 23^, although several key features distinguish Aer from receptors with periplasmic ligand binding domains. An asymmetric “piston motion” of the TM2 helix in the canonical aspartate receptor Tar is likely a conserved feature in many receptors^24–26^. The methylation helix cap (MH-cap, made up of a dimer of N-terminal MH1 and C-terminal MH2 helices) is an antiparallel 4-helix bundle that intersperses the HAMP and the methylation helices of canonical MCPs^27, 28^. In most MCPs, the methylation helices contain conserved Glu residues that undergo reversible methylation by the CheR/CheB methyl transferase/methyl esterase system to tune receptor output to kinase activity^10, 29^. An evolutionarily conserved “LLF” motif at the C-terminal end of MH2 couples the HAMP to the KCD and controls HAMP-mediated receptor signal output in the *E. coli* serine receptor Tsr^30^. In Aer, the TM region is thought to have no direct signal transduction role but may function to integrate Aer into the chemosensory arrays and juxtapose the PAS domains relative to other membrane components^31^. In addition, despite possessing conserved Glu/Gln residues that typically serve as methylation sites in the KCD, Aer appears not to undergo adaptation-related methylation^15^. Thus, we will refer to the MH-cap region in Aer as a “methylation helix-like cap” (MHL-cap). Differences at the LLF motif in Aer (VLL) also suggest a different proximal signaling mechanism from conventional MCPs.

The PAS domain of Aer binds a redox-sensitive flavin adenine dinucleotide (FAD) cofactor^6, 16, 17^. As a flavoprotein, Aer stabilizes the reduced anionic semiquinone (ASQ) state of FAD^32, 33^, as shown by *in-cell* ESR spectroscopy, which revealed the predominance of the ASQ radical state *in vivo*^34^. Changes in FAD redox state induce conformational changes that propagate down to the KCD, which relays signal to CheA and the chemotaxis signaling cascade^4, 16, 17^. Oxidation of Aer-FAD to the quinone state promotes a kinase-on state, whereas reduction to ASQ under hypoxic conditions inhibits CheA^32^. Physiological data indicates that a fully reduced hydroquinone (HQ) state of Aer induces CheA activation as well^35^. However, the HQ state is difficult to generate in purified Aer^33^, suggesting that a fully reduced flavin signal may involve an apo form of the protein. A currently unidentified species associated with the ETC modulates the redox state of Aer^35^. NADH dehydrogenase activity correlates with Aer-mediated motility and could serve as an electron donor, especially because the reduction potential of oxidized Aer is quite low at < -290 mV^33^. Known structures of FAD-binding PAS domains include the NifL PAS domain from *Azotobacter vinelandi* and many members of the LOV-family of photoreceptors, such as AsLOV2 and Vivid from *Neurospora crassa*^36–38^. In these cases, protonation of the flavin N5 atom in concert with flavin reduction or cysteinyl adduct formation initiates signal propagation, altering the hydrogen bonding network within the FAD-binding pocket. For the LOV proteins, a Gln residue on the Iβ strand responds to changes in the flavin redox and protonation state. In Aer this residue is an Arg (115) and the crystal structure of an Aer-PAS GVV variant (in which three substitutions were made to enhance stability of the isolated domain: S28G, A65V, and A99V) shows that Arg115 resides beside the flavin N5 but is just outside hydrogen bond distance^33^.

HAMP domains of canonical MCPs act as the central processing units and transmit signals “vertically” from the TM region^11, 39^. Aer HAMP bears strong sequence similarity to these HAMP domains but its role in signal transduction may be mechanistically different because of the “lateral” signal transmission from the PAS domains. Spin-labeling studies demonstrate that the PAS domains associate closely with the HAMP both in vitro and in vivo^32, 34^. Mutations promoting a signal-on state of Aer cluster in the PAS-HAMP interface, suggesting a signalling role for these domain contacts^40, 41^. The parallel four-helix bundle of the HAMP comprises a dimer made up of subunits each with two amphipathic α-helices (AS-1 and AS-2) bridged by a connector loop. FAD binding and aerotactic activity is lost upon mutation of the AS-2 helix in the HAMP domain, but is rescued by the PAS-GVV mutant^33, 42^. PAS domain stability relies on residues in the HAMP domain, presumably through their shared interfacial interactions. Cysteine substitution crosslinking and solvent accessibility studies confirm close proximity of the PAS β-scaffold and AS-2 HAMP helix^40, 41^. A range of models have explored HAMP signaling^11, 26, 43–49^, with elements of static-dynamic transitions and helical rotations supported by structural and biochemical data. In general, more structural data is needed to understand whether HAMP domains convert disparate conformational inputs (i.e. canonical MCP TM signal transduction, lateral PAS signal transduction) into similar outputs, or play different roles in different receptors.

Here, we use cryo-electron microscopy (cryo-EM) single-particle analysis (SPA) to report the structure of full-length Aer in its oxidized (Q) and reduced (ASQ) states. UV-Vis spectroscopy confirms FAD reconstitution and reduction of solubilized Aer to the ASQ. Biochemical analyses demonstrate that the detergent-solubilized receptor modulates CheA in a redox-dependent manner. Conformational changes at the FAD-binding pocket and PAS-MH interface are observed upon Aer reduction that dramatically alter the stability of the KCD. With spin-labeling and ESR spectroscopy, we further probe the structural and dynamic changes of the KCD in kinase-on and kinase-off states. Taken together with existing data, a consensus conformational signaling mechanism for the initial stages of Aer conformational signaling emerges.

## Results

### Aer purification, activity and redox states

For structural characterization of active WT Aer by cryo-EM, we developed a lauryl maltose neopentyl glycol (LMNG)-based solubilization and purification workflow from detergent screens of the protein expressed in *E. coli* BL21 cells (SI Appendix, Fig. S1A). Co-expression with chemotaxis proteins CheA and CheW substantially improved Aer yield. The 150 kDa homodimer of WT Aer solubilized in 1% LMNG retained FAD-binding and regulatory function of CheA autophosphorylation (SI Appendix, Fig. S1B, Fig. S2). Dithionite reduces WT Aer to the anionic semiquinone (ASQ) state in anaerobic conditions, inducing a kinase-off state. UV-visible and ESR spectroscopy verified the redox state of the cofactor in the reduced protein (SI Appendix, Fig. S2AB). ASQ Aer solubilized in LMNG inhibits CheA autophosphorylation activity (SI Appendix, Fig. S2C). Further reduction to the hydroquinone (HQ) state was not observed and Aer retained FAD upon reduction to ASQ and reoxidation (SI Appendix, Fig. S2D-F). Purified Aer was applied to cryo-EM grids in oxidized or reduced states, vitrified, and imaged to obtain the structure of Aer in both the oxidized flavin quinone form (Q) and the ASQ form (SI Appendix, Fig. S3, Table S1). 2D and 3D classifications of the particle stack for Aer Q revealed two separate populations of conformations: MHL-free and MHL-bound, referring to the interaction of the PAS sensory module with the proximal methylation helix cap of the KCD (Fig. 2, SI Appendix, Fig. S3A, S4). The MHL-cap does not interact with the sensory module of Aer in MHL-free and the density of the KCD is unresolved owing to high conformational heterogenetity (Fig. 2B). However, for MHL-bound particles, we observe the stabilization and visualization of the membrane-proximal region of the KCD through interactions between the PAS domains and MHL-cap (Fig. 2B, SI Appendix, Fig. S4AB). Both populations were refined to yield a global resolution of 3.5 Å (90k particles) and 4.0 Å (14k particles) respectively (SI Appendix, Fig. S3A,Table S1). An estimated 10% of the oxidized particle set adopted the MHL-bound conformation, suggesting that incorporation of single receptors in micelles favor the more dynamic state (SI Appendix, Fig. S3). We first describe the structural conformation of oxidized Aer according to the higher resolution structure corresponding to the more prevalent MHL-free oxidized state.

### The structure of oxidized Aer Q MHL-free

Aer Q forms a C2 symmetric homodimer, dimerizing through the TM region, HAMP module, and KCD (Fig. 2A,B). Whereas we observe dimerization in TM and HAMP, we only observe KCD dimerization in the MHL-bound form. The HAMP-proximal PAS domains bind the FAD cofactor deep in the PAS pocket, with the adenosine moiety exposed on the surface (Fig. 2C). The LMNG micelle encapsulates the TM region, with the F1-linker situated between the membrane and the PAS domain. The positively charged and hydrophobic F1-linker region (Fig. 2D) important for FAD-binding^6^ positions the PAS domain through interaction with the membrane surface (Fig. 2D). F1-linker residue W152 inserts directly into the micelle (Fig. 2E).

The PAS domain conformation resembles the previously determined PAS-GVV structure (PDB ID: 8DIK)^33^: five antiparallel β-strands and peripheral α-helices assemble to form the PAS core. An N-terminal coil flanks the core (Fig. 2E)^33^. Removal of N-cap residues has been linked to conformational changes in Aer that mimic a signal-on state of the receptor^50^. Similarly, signal-on mutations in Aer cluster at the N-cap^40^. Residues 1-5 of the N-cap do not associate with the PAS domain and are too flexible to capture in the reconstruction (Fig. 2B).

FAD is nestled in the binding pocket of the PAS core, as shown in the PAS GVV structure (SI Appendix, Fig. S5A). However, in contrast to the PAS-GVV conformation, in MHL-free Aer, the pocket residue R115 hydrogen bonds to FAD N5 and O4 in the oxidized state. The guanidine group of R115 also closely interacts with S22 and H53, stabilizing the FAD position (Fig. 2C). Unlike PAS-GVV, where N85 hydrogen bonds with both FAD N3 and O4, in MHL-free Aer (and more so in MHL-bound Aer) N85 shifts to only hydrogen bond with FAD N3. The FAD moiety resides further out of the PAS pocket than in the PAS GVV (flavin N5 to R115 N distances of 8.8 Å to 8.2 Å respectively), as reflected by the relative positioning of the side chains of R115 and H53 (SI Appendix, Fig. S5A). F37 has also flipped relative to its position in PAS-GVV. Without the GVV substitutions that increase stabilizing interactions with the isoalloxazine ring, isolated Aer PAS does not bind FAD^33^, likely owing to loss of the HAMP AS-1 helix that serves as a C-cap against the PAS β-sheet (Fig. 2E), as is found in other PAS domains.

The HAMP domain of Aer folds into a canonical parallel four-helix bundle composed of the AS-1 helix that follows from the transmembrane region, a connector region, and the AS-2 helix that connects to the KCD region(Fig. 2E). Alterations in the helical heptad repeat packing of HAMP domains, particularly due to changes in helix rotation, have been associated with the propagation of conformational signals in canonical chemoreceptors. For example, changes in helical positioning that agree well with transitions between “x-da” to more typical “knobs-into-holes packing” have been associated with signal transduction in the Tsr serine receptor and others^28, 43, 44, 46, 51–55^. Three of the four hydrophobic packing layers of the Aer HAMP shows mainly knobs-into-holes packing, although the second layer involving T242, V215, and A216, has the “x-da” type geometry (SI Appendix, Fig. S6A). Notably, the relative positioning of the AS-1 and AS-2 helices are shifted by half a helical turn compared to previously determined structures of HAMP domains. Structural homology searches^56^ reveal that the HAMP helical packing arrangement resembles that of some HEAT repeat proteins (SI Appendix, Fig. S6B). Helices AS-1 and AS-2 are linked by a connector that contains an internal salt bridge between R227 and E231 (Fig. 2E), and provides key interactions to the PAS domain through G225, E226, Q248, and H232 and to the F1 region through S236 and D237 (Fig. 3A). D237 belongs to the conserved DExG motif (residues 237-240) at the top of AS-2^11^, where E238 reaches across to interact with E213 of AS-1 (Fig. 2E, 3A). AS-2 binds the PAS β-sheet and provides several sidechain contacts from R244, Q248, L251 and R254, whereas AS-1 provides only minimal contacts on the adjacent subunit at the transition to the connector. C253 and its symmetry mate have their thiol groups within 7 Å but do not form a disulfide, although a disulfide has been observed when treated with an oxidant^57^. Sequence analysis revealed a phase shift in the heptad registry of the AS-2 helix at residues 254-256 of the HAMP-KCD connection^11^, and was further refined to be between Aer residues 259-262 by cysteine modification studies^41^. Here, the resolvability of the map in the MHL-free class drops in this region with no clear density for residues 254 and onwards (SI Appendix, Fig. S5B).

**Figure 3:**
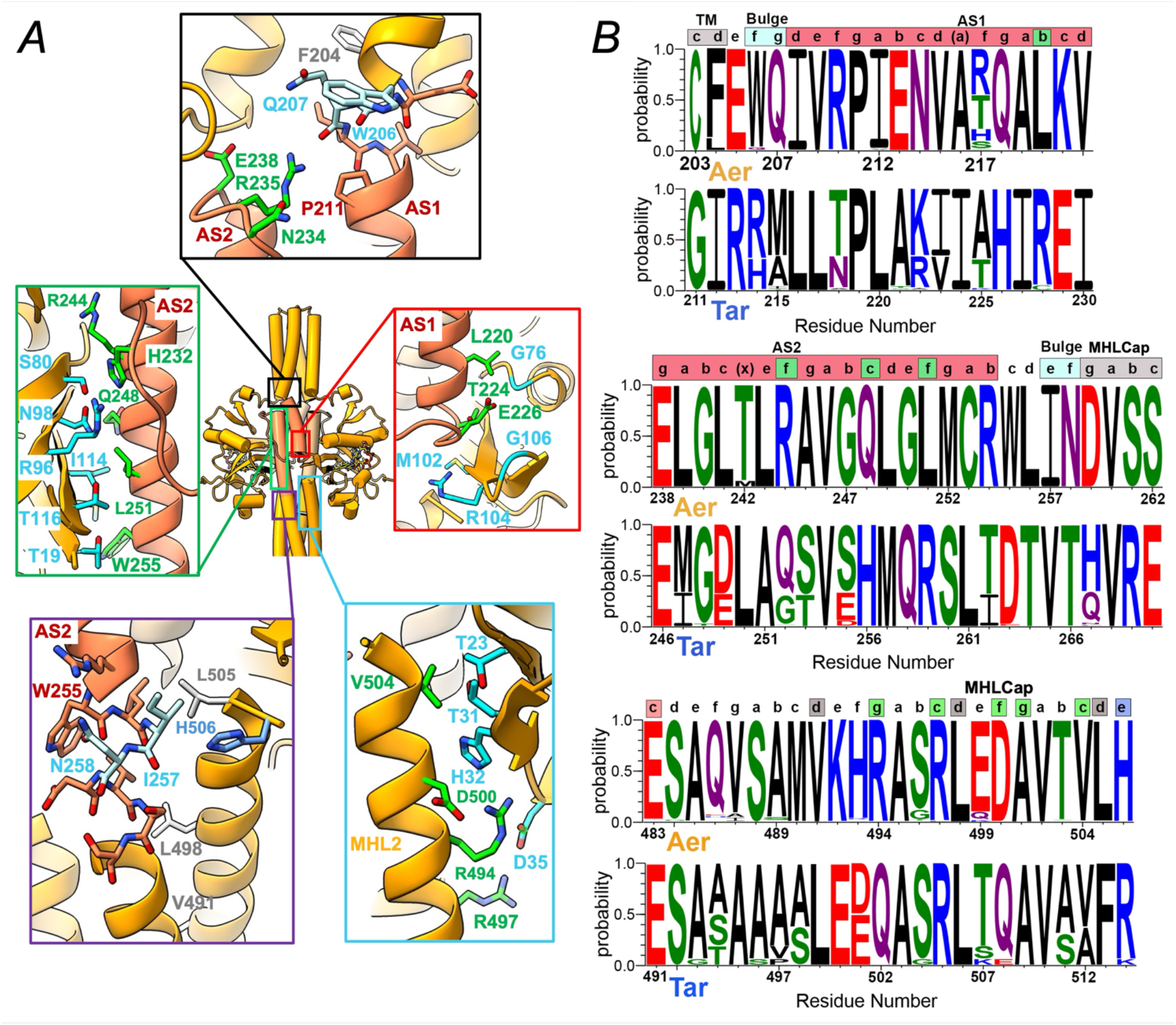
Aer HAMP domain interactions and domain coupling. (A) The HAMP domain of Aer Q MHL-bound (coral ribbons) is offset from the transmembrane region by a phase shift in heptad packing that creates a bulge at W206/Q207 (cyan bonds, top black-boxed inset). P211 on helix AS1 prevents interhelical hydrogen bonding, which is compensated by R235 (green) on the connector interacting with the bulge peptide backbone. HAMP residues (green) from AS-1 (red boxed inset) and connector and AS-2 (green boxed inset) stabilize interactions with the flanking PAS residues (cyan), which are protected in the interface from crosslinking agents^41^. The HAMP domain is offset from MHL1 in the MHL-cap by a phase stammer that creates a bulge at I257, N258 (cyan), which is stabilized by main-chain hydrogen bonding from H506 on MHL2 (purple boxed inset). Residues on MHL2 (green) interact with PAS domain residues (cyan) in the Cα, Dα region (cyan boxed inset). V491, L498 and L505 substitute for a LLF motif that stabilizes a zipped helix cap structure in Tsr. (B) Sequence logos for Aer receptors (above) and Tar receptors (below) for the TM-to-AS1 transition (top), the AS2-to-MHL1 transition (middle), and the MHL2 C-terminus (bottom). The top matching 500 non-redundant Tar and 250 non-redundant Aer sequences were aligned and compared. Heptad positions assigned by Socket2 are shown above and colored according to their residue depiction in (A) and (C). Sequence logos are colored with respect to the chemical identity of the residue (positive – blue, negative – red, hydrophobic – black, hydrophilic – purple, C, G and T – green).

In chemoreceptors that contain LBDs, the TM region is responsible for transmission of signal from the periplasmic sensing domain to the cytoplasm, a key feature of which has been described as a piston-like motion of TM2^10, 24^. TM2 conformational changes are transmitted to the HAMP domain through a five-residue control cable segment^58^. Of these, “trigger” residue I214 plays a critical role in Tsr TM-HAMP signaling^58^. Because Aer is not known to transmit signals through the membrane it lacks classical control cable input and the TM-HAMP signaling function of this region is lost. The analogous control cable of Aer consists of CFEWQ, with F204 replacing the isoleucine trigger residue (Fig. 2E, 3). As part of the hydrophobic core of the TM helix packing, the rigid aromatic ring of F204 likely reduces conformational flexibility at the end of the TM region (Fig. 2E, 3A). The LMNG micelle largely buries the periplasmic loop between TM1 and TM2 (residues 183-187) despite some of these sites being accessible to membrane impermeable labeling agents^31^.

### The structure of oxidized Aer Q MHL-bound

Compared to the more prevalent MHL-free conformation, we observe a compression along the axis parallel to the membrane in the MHL-bound state of Aer (Fig. 4A). The distance between the N5 atoms of the FAD molecules is reduced by approximately 3 Å (Fig. 4A) upon PAS-MH interaction. TM, PAS-HAMP, and HAMP-HAMP interfaces rearrange to bring about the compacted state (Fig. 4AB). For MHL-bound, a constriction of the interface between the PAS and HAMP domains (Fig. 4A) offsets relaxed HAMP-HAMP subunit contacts: i.e. the interchain distance between equivalent AS-1 and AS-2 helices increases nearly 2 Å (Fig. 4B). The expansion of the HAMP-HAMP module may indicate a decreased stability of the HAMP region, conversely corresponding to an increased stability of the KCD. Studies of chemoreceptors suggest that HAMP-HAMP stability is inversely correlated with KCD stability^28, 51^. Residues previously identified in the PAS-HAMP interface (N98 (Hβ), I114 (Iβ), and Q248 AS-2)) (Fig. 3A) have altered interactions in the compact and expanded states^40^. Previous data shows that PAS mutation N34D acts as an allele-specific suppresor of HAMP mutation C253R, which resides in the HAMP-HAMP interface at the base of AS-2 (Fig. 4C)^42^; hence in this double mutant, modest destabilization of PAS-HAMP may be compensated by even greater destabilization of HAMP-HAMP.

**Figure 4:**
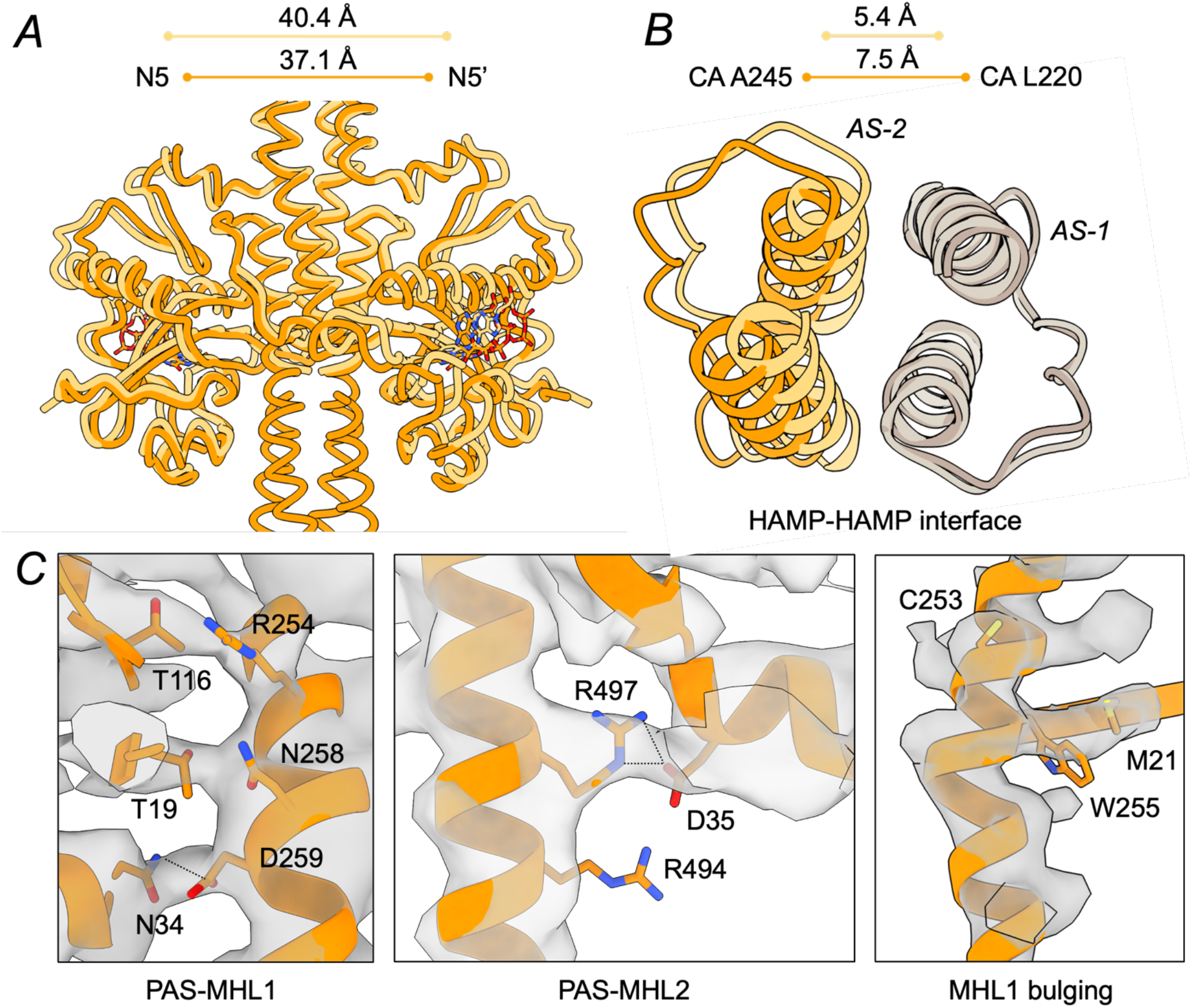
Oxidized Aer compresses laterally upon binding the MHL-cap of the KCD. (A) Cartoon representation of MHL-free (tan) and MHL-bound (orange) conformations overlayed, showing overall compression of the MHL-bound structure compared to MHL-free (FAD N5 to N5’ distance: 40.4 Å for MHL-free, 37.1 Å for MHL-bound). (B) The dimeric HAMP-HAMP interface relaxes upon binding MHL. A245 CA on AS-2 to L220 CA on AS-1 distance: 5.4 Å for MHL-free, 7.5 Å for MHL-bound. (C) Interacting residues of PAS-MHL1 and PAS-MHL2 (hydrogen bonds, dotted lines) in MHL-bound. The MHL-bound structure shows bulging of the MHL1 helix at the PAS-MHL1 interaction interface (see Fig. 3), opposite of the M21-W255 interaction.

PAS-MHL-cap interactions are mediated by proximal residues on both MHL1 and MHL2 (Fig. 3A, 4C). Both KCD helices interact with PAS Aβ, Bβ and Cα through packing and electrostatic attractions, the latter involving N34 (Cα) with D259 (MHL1), and D35 (Cα) with R494 and R497 (MHL2) (Fig. 3A, 4C). Substitutions of N34 severely affect receptor activity, often initiating a signal-on bias^19, 40^.

The parallel helix packing HAMP domain is offset from the antiparallel helix packing TM and MHL-cap regions by two phase shifts in the helical heptad repeat (Fig. 3B). Structurally, these discontinuities both manifest as bulges/kinks in the helices that join TM2 to AS-1 and AS-2 to MHL1. At the TM2-AS-1 junction, a residue insertion at W206/Q207 bulges the helix such that these residues do not make interhelical i to i+4 hydrogen bonds (Fig. 3A). The bulge coincides with the position of P211, which cannot supply an amide nitrogen, and R235 on the connector that hydrogen bonds to the Q207 main-chain. Similarly, the structural distortion at the AS-2-MHL1 junction involves an insertion at two non-helical hydrogen bonding residues in L257-N258 and the main chain carbonyl of I257 accepting hydrogen bonds from the Ser 261 sidechain and the C-terminal residue H506 on MHL2, which have similar roles as P211 and R235 in TM2-AS-1 junction (Fig. 3A). Directly N-terminal of the AS-2-MHL1 bulge, W255 packs against M21 of PAS Aβ, an interaction absent in the MHL-free structure (Fig. 3A, 4C). C-terminal to the bulge, the MHL1 helix shifts away from the longitudinal axis of the receptor stalk and reduces hydrophobic packing with the adjacent subunit. From cysteine modification studies, the phase shift at the AS-2-MHL1 junction was defined at positions 259-262^41^; this is indeed the position of discontinuity in the heptad packing arrangement, which is best described as a phase “stammer” whereby the heptad repeat offsets by a deletion of 3 positions, transforming an “a” to a “d” position (see Fig. 8 in^46^, and SI Appendix, Fig. S6C). However, this transition in helical packing is facilitated by the bulge at residues 257-258 (Fig. 4C).

### The structure of reduced Aer

LMNG-solublized Aer was reduced to the ASQ state through treatment with dithionite (SI Appendix, Fig. S2) and subsequently vitrified under anaerobic conditions, and then imaged to obtain a structure with a global resolution of 3.3 Å (SI Appendix, Fig. S3B). The structure of the reduced Aer ASQ state strongly resembles the oxidized MHL-free population (peptide backbone root mean square deviation of 0.8 Å, Fig. 5A,B). Both structures can be resolved up until the phase shift at the C-terminus of AS-2 (Fig. 5C, SI Appendix, Fig. S5B, S7B).

**Figure 5:**
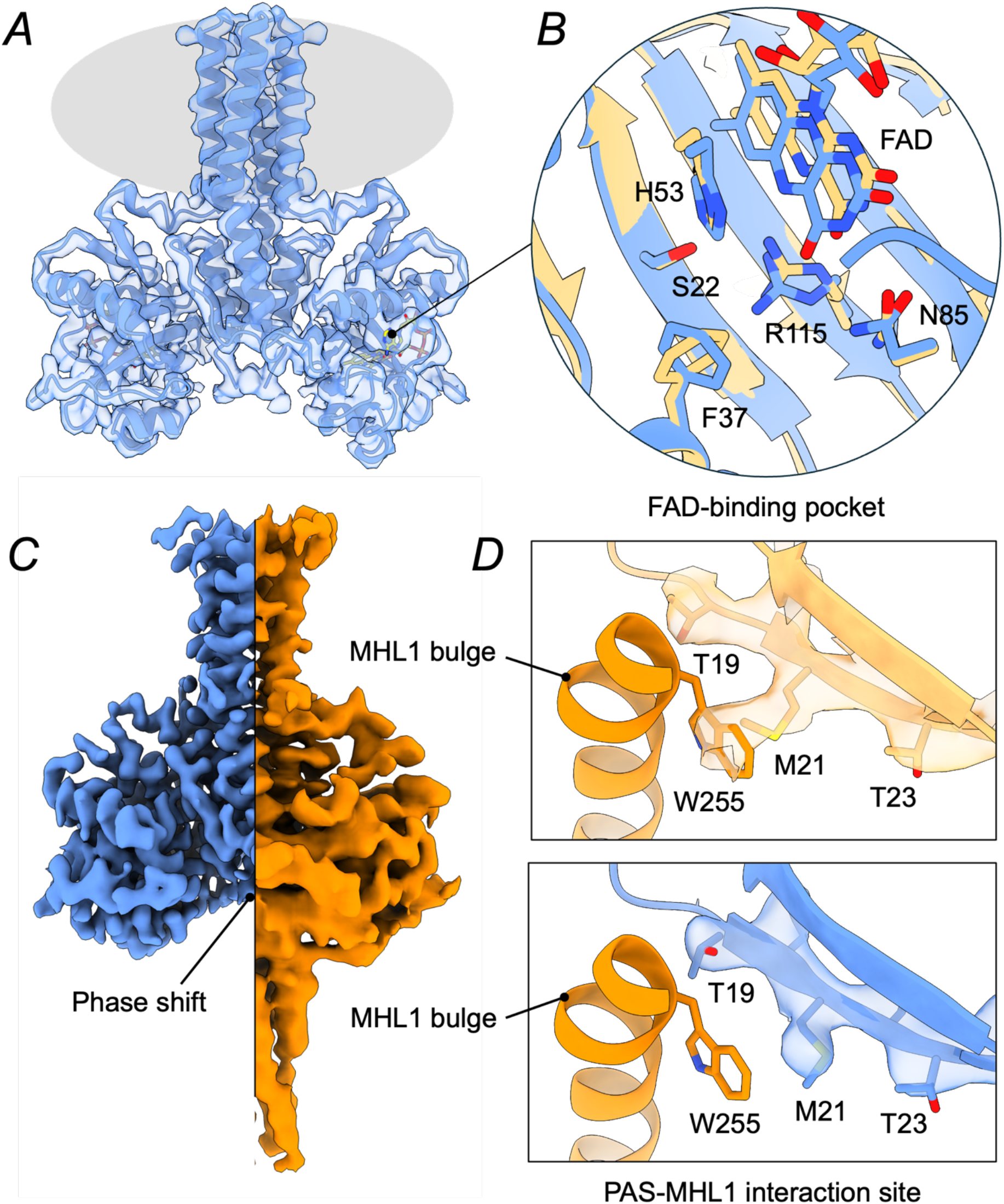
Reduced ASQ Aer resembles an MHL-free conformation. (A) Cartoon representation of the structure of ASQ Aer (blue) fit into its electron density map (LMNG micelle area, gray). (B) Detailed view of the FAD-binding pocket, with the structure of MHL-free superposed onto ASQ Aer. (C) Electron density comparison of ASQ (blue) and MHL-bound Q (orange) Aer, showing the lack of KCD density for ASQ. (D) Detailed view of ASQ (blue) and MHL-free Q (tan) Aer PAS-MHL interaction shown with electron density. Positioning of M21 changes between the MHL-free Q state (top, tan) and the ASQ state (bottom, blue). The MHL1 helix in an MHL-bound Q (orange) state is incompatible with M21 in MHL-free Q Aer, indicating that redox-induced changes in the PAS-domain can affect MHL-cap interactions.

Although the overall structures of Aer Q MHL-free and Aer ASQ are similar, there are distinct differences in the FAD binding pocket and the PAS-MHL-cap interface (Fig. 5B, SI Appendix, Fig. S7A). In Aer ASQ, R115 no longer interacts with N5 of FAD and instead forms hydrogen bonds with S22 and S113 (Fig. 5B, SI Appendix, Fig. S7A). In response, the F37 aromatic ring flips to form a π-cation interaction with R115, similar to that found in PAS-GVV (SI Appendix, Fig. S5A). The positioning of FAD in the binding pocket also shifts to be more PAS-GVV like, with the isoalloxazine ring sinking further into the cleft in the reduced structure (flavin N5 to R115 N distance of 8.1 Å). N85 also now hydrogen bonds to both flavin N5 and O4. (The correspondence of PAS-GVV to Aer ASQ raises the possibility that the Aer GVV crystals were at least partially reduced in the X-ray beam). The structural changes in the Aer ASQ flavin pocket couple to a compression of Aβ and movement of the M21 side chain that packs against W255 in Aer Q MHL-bound (Fig. 5D, SI Appendix, S7A). As a result of such altered PAS-MHL interactions (Fig. 5D), the KCD is likely more dynamic and conformationally heterogeneous.

The role of R115 in Aer-PAS redox-signaling resembles that of the conserved Iβ Gln residue of light-sensitive LOV domains (SI Appendix, Fig. S7C). The hydrogen bonding network of Q182 changes upon protonation of the flavin ring at N5, correlating with movement of both N-cap and C-cap restructuring, depending on the LOV protein^59^. Whereas any N-cap alteration is beyond detection at the resolution of the structures, the MHL-interacting PAS β-strands appear to respond upon R115 repositioning.

The Aer ASQ structure also has a compressed HAMP interaction and expanded PAS-HAMP interaction compared to Aer Q MHL-bound (SI Appendix, Fig. S8A). The compressed HAMP packing is also somewhat reflected in the TM region, wherein the F204 residues rotate to make closer contacts in the reduced state, perhaps contributing to enhanced stability of the TM-HAMP unit (SI Appendix, Fig. S8B). Both in vitro and in vivo pulse-ESR measurements of the Aer ASQ state correspond to the spacing measured in both the MHL-free and reduced structures^32, 34^. In the ASQ state, PAS-HAMP interactions relax and the distance between the PAS domains increases; concurrently the HAMP-HAMP packing increases (SI Appendix, Fig. S8A), perhaps accentuating the instability in the KCD packing resulting from the helix bulge at the phase shift. Cysteine substitutions of W255 and L256 at the MHL1 bulge disrupt Aer-mediated aerotaxis, but not protein stability, emphasizing the role of these residues in signalling^60^. The combined effects of PAS-MHL-cap destabilization and tighter HAMP packing likely favor the MHL-free bound configuration of the KCD. 2D classification of the Aer Q and ASQ data sets show significant differences in KCD density (SI Appendix, Fig. S4A). For oxidized Aer Q, the KCD structure is static enough to allow alignment of the coiled-coil helices in some of the 2D classes, whereas none of the reduced conformations show any density beyond the phase shift (SI Appendix, Fig. S4). Distal interactions at the PIR are likely affected by the change in KCD dynamics, indicating a PAS-to-MHL-cap dependent signaling mechanism.

### KCD properties from DEER spectroscopy

To examine KCD conformational dynamics in full-length Aer receptors, we incorporated the proteins into nanodiscs to mimic a phospholipid bilayer (SI Appendix, Fig. S9) and applied double electron-electron resonance (DEER) spectroscopy to nitroxide (MTSL) spin-labeled positions on the KCD for both the oxidized Q state and a proxy of the reduced ASQ state, a D68V kinase-off variant (Fig. 6A). The nitroxide radicals that report on the protein conformation are sensitive to reductant and thus, Aer under reducing conditions could not be directly tested. The D68V substitution, which resides on PAS αF, was identified as constitutive kinase-off mutant^40^. Aer D68V incorporated into nanodiscs substantially reduced CheA autophosphorylation and subsequent phosphotransfer to CheY relative to that of Aer WT (SI Appendix, Fig. S9BC), findings consistent with the smooth swimming behavior observed in vivo^40^. Eleven single cysteine sites were engineered into Aer with the two native Cys residues substituted to Ser (Δcys) with the intention of measuring distributions of separation across the dimer interface (Fig. 6A): two were placed in the HAMP domains at the C-termini of AS-1 and AS-2, and nine sites were introduced into the KCD. Eight of these KCD positions were chosen based on molecular modelling predictions to report on the structure and dynamics of the N- and C-terminal α-helices at specific locations along the four-helix bundle. P(r) distributions constructed from 4-pulse-DEER data on each of the variants generally exhibited two populations, a shorter-distance population with a peak maximum between 30-34 Å and another, longer-distance population with a peak maximum between 43-50 Å (Fig. 6B, SI Appendix, S10). Comparisons of P(r) distributions of MTSL-labelled Aer mutants in nanodiscs prepared from separate purifications were made to assess reasonable deviations in shorter- and longer-distance peaks and to evaluate the reproducibility of the relative peak populations from sample to sample (SI Appendix, Fig. S10). The maxima of the shorter-distance peaks differed by only 1.4 Å and when fit to gaussian functions, the standard deviations of the peaks (σ) differed by 0.36 Å. The average shorter-distance peak (calculated for all samples) of 32 ± 3 Å reflects the separation between spin labels located at symmetric sites of a chemoreceptor dimer^32, 52, 54, 55^. Shorter distance peaks in the P(r) distributions were fit to Gaussian functions for additional analysis.

**Figure 6:**
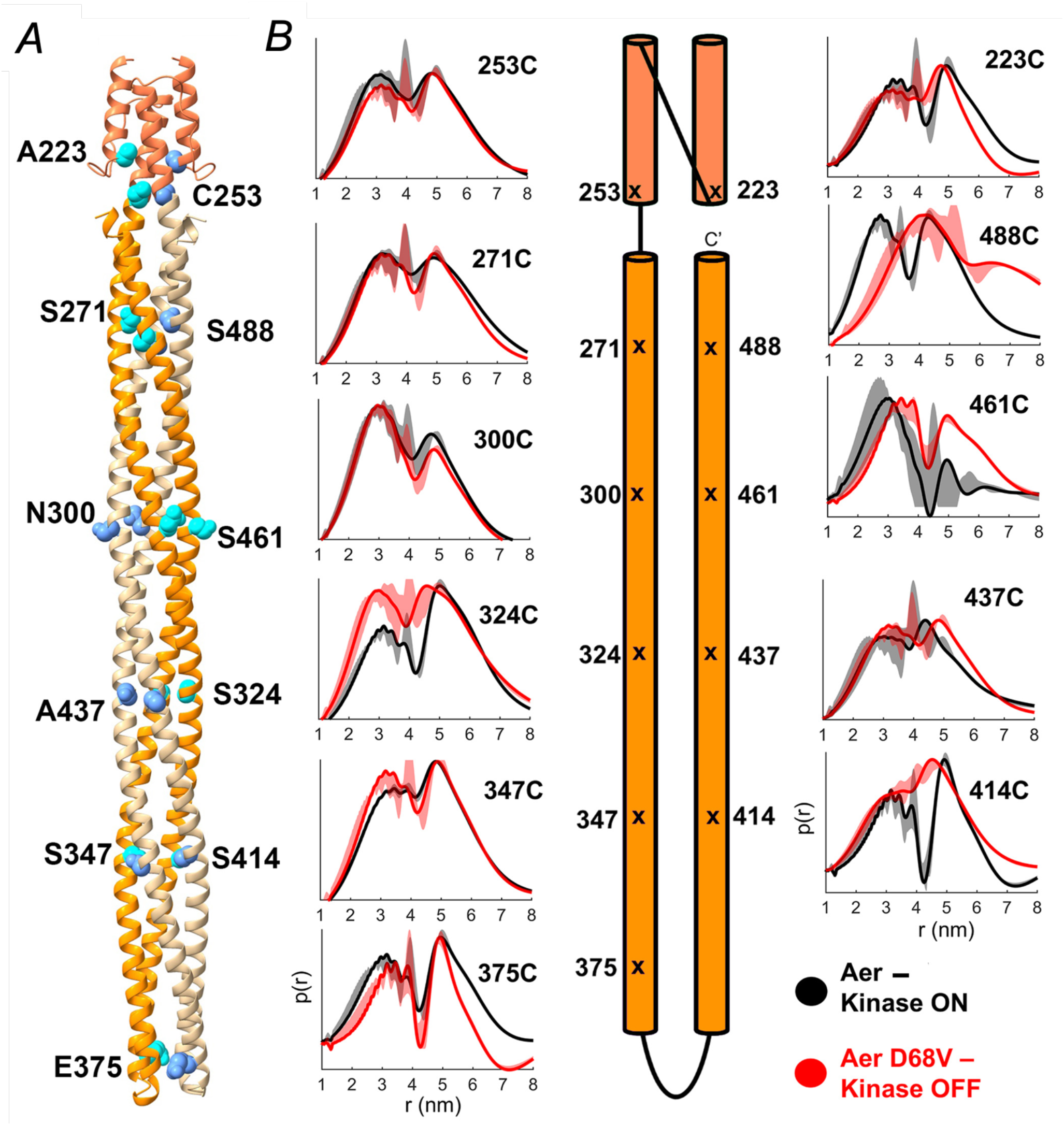
Subunit-to-subunit distance distributions for spin-labeled Aer KCD sites. (A) AlphaFold 2 model of Aer showing the HAMP (coral) and intact KCD domains (orange and tan for each subunit), with the spin-labeling sites depicted as space-filled sidechains (cyan). (B) P(r) distributions for each site along the N-terminal helix (left) and the C-terminal helix (right) separated by a schematic representation of the spin-labeling placement. Aer Δcys + label representing kinase-on is in gray; Aer Δcys D68V + label in red. The distance domain was obtained by using the SF-SVD method. Errors in the distance distributions are represented by gray and red shading. Time domain data shown in Figure S10.

The population and maxima of longer-distance peaks were significantly less reproducible. Several factors may contribute to the inconsistency of the longer-distance signal observed in the P(r) reconstructions. 1) Longer distances may represent PDS signal from static disordered states, 2) the longer-distance peaks may represent PDS signal from two crosslinked Aer dimers, which cannot be completely avoided during spin-labeling. This possibility may also account for the relatively broad and inconsistent features of the longer-distance peaks, as additional states are accessible to the crosslinked species through rotations and conformational changes across the disulfide bond.

### Conformational differences in the KCD between signaling states

Comparisons of the DEER separation distributions at specifically labelled residues in Aer Δcys and Aer Δcys D68V offer insights into the structural and dynamic changes that occur upon signal switching local to that residue. The P(r) distributions report on the distances between the same residue of each Aer subunit within the membrane-embedded dimer. Secondly, the average distance of the P(r) distribution below 40 Å, <R>_<40Å_, was compared to assess changes in the near, more accurate separation distributions (Fig. 6, SI Appendix, Table S2). Two residues in the HAMP domain at the N-termini of AS-1 (A223C) and AS-2 (native C253) were labelled, as well as nine residues in the KCD, and one (E375C) in the PIR.

Consistent with the cryo-EM results, the most dramatic changes in A68V compared to Aer Δcys occured at the C-terminal region of MHL2 in sites 461 and 488 (Fig. 6B). At position 461, the distance peak of Aer Δcys D68V (34.3 Å) expands compared to that of Aer Δcys (30.5 Å). Position 488, closest to the PAS-MHL-cap interface reveals the most dramatic change, with the shorter distance distribution shifting 15.2 Å further than in Aer Δcys D68V. Furthermore, <R>_<40Å_ differs between the two samples by 4.1 Å, indicating a much more extended ensemble (SI Appendix, Table S2). This result indicates that the C-terminal regions of MHL2 separate by approximately 7.5 Å and obtain much greater flexibility. This dramatic change may indicate liberation of this region of the C-terminal KCD helix from contacts with the helix bundle altogether.

In contrast, all other probed positions display only modest changes to P(r) between signaling states (Fig. 6B, SI Appendix, Table S2). Although no one position exhibits major differences in the distance distributions between the kinase-on and kinase-off signaling states, the minor changes that are observed suggest a pattern. In general, probed residues along the N-terminal α-helix exhibit either no significant changes in distance distribution or modestly shorter distances in Aer Δcys D68V samples than in Aer Δcys. Conversely, residues along the C-terminal α-helix tend to exhibit longer distances in Aer Δcys D68V relative to Aer Δcys.

## Discussion

The Aer structures are informative with respect to the extensive genetic and biochemical analyses previously performed on the system. Mutations that cause kinase-on (Fig. 7A,B) and kinase-off (Fig. 7C,D) biases in Aer map to regions of the structure consistent with a PAS-to-MHL-cap signaling axis, but also reflect the coupled nature of entire HAMP-PAS-MHL-cap unit^19, 40^. That said, kinase-on substitutions predominantly cluster around the flavin pocket; many of them would be predicted to be detrimental to flavin binding (Fig. 7AB). Thus, disruption of FAD binding in the PAS domain may deliver a kinase-on signal. Aer mutants that retained FAD binding but abolished aerotaxis are concentrated at the region of the AS-2-MHL1 bulge (Fig. 7A), highlighting the functional role of this region. Accessibility of Aer to PEGylation probed by cysteine replacement mutagenesis indicate that kinase-off states greatly increase accessibility of the MHL-cap relative to kinase-on states, consistent with both the cryo-EM and DEER data (Fig. 7E)^41^. Kinase-off states also decrease accessibility of HAMP residues to PEG within the hydrophobic core, consistent with the observed contraction of the HAMP bundle in Aer ASQ. In contrast, PEG accessibility of HAMP Cys replacements in the PAS-HAMP interface increase in the on-state, counter to the tighter PAS-HAMP interface found in cryo-EM for Aer Q. It is difficult to reconcile these findings completely, although the loosened HAMP-HAMP contacts in the kinase-off states may favor larger scale processes, like overall HAMP conformational fluctuations or subunit exchange that could increase overall accessibility during the relatively long time scales of the PEGylation reactions. It is also possible that the Cys substitutions discourage a tight PAS-HAMP interaction.

**Figure 7:**
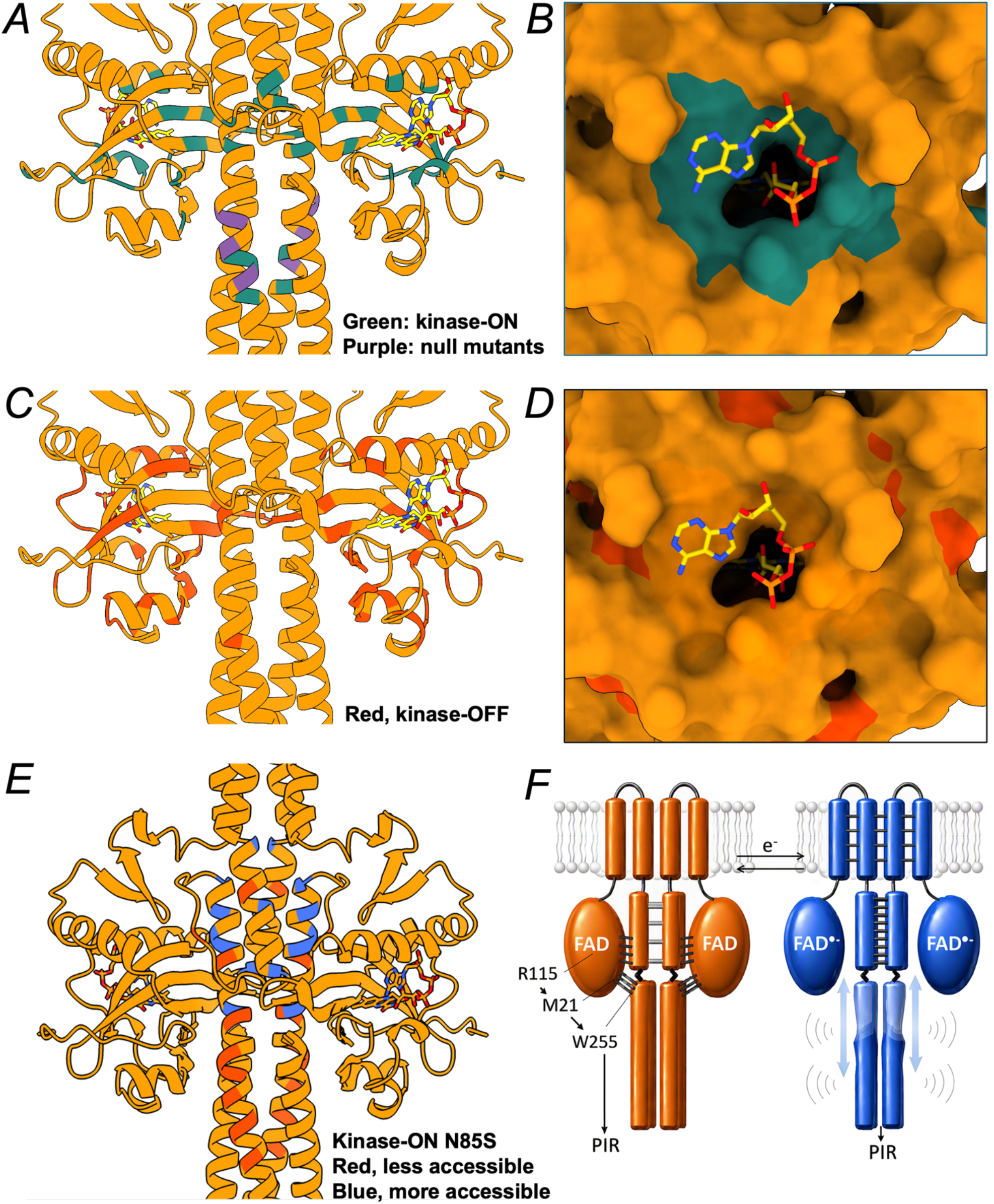
Mapping Aer mutations of known function onto the MHL-bound structure. (A) Sites of kinase-on mutations (green) and mutations that abolished aerotaxis but retained FAD-binding (purple)^19, 40, 74^. (B) Surface representation of the FAD-binding pocket of Aer, showing the clustering of kinase-on mutation sites (green) around FAD (yellow)^19, 42^. (C) Sites of kinase-off mutations (red) and (D) their distal clustering with respect to the FAD binding pocket^19, 42^. (E) Changes in accessibility to PEGylation of engineered cysteine substitutions in a kinase-on variant (N85S) compared to WT^43^. Red positions were less accessible in kinase-on, blue sites more accessible. (F) Schematic summarizing changes in Aer conformation upon reduction to the ASQ. Flavin reduction rearranges R115 and other coupled residues in the flavin pocket. These changes affect the PAS β-strands and alter the conformation of M21, which in turn affects the PAS-MH-cap interface, particularly interacting residue W255. Weakening of PAS-HAMP interactions also strengthens HAMP-to-HAMP subunit contacts. Both effects likely conformationally destabilize the MHL-cap region of the KCD, particularly MHL2, the C-terminal helix. These changes in helix dynamics propagate to the PIR.

In Aer, signal transduction is initiated when reduction of the flavin cofactor to the ASQ state alters the charge distribution in the active center, notably increasing electron density at N5 (Fig. 7F). This shift reorganizes R115 and its interacting residues within the flavin pocket, producing a modest but globally distributed PAS-domain response, especially in β-strands whose distal residues modulate contacts with the MH-cap helices—most prominently the Met21–Trp255 interaction (Fig. 7F). Concomitantly, PAS–HAMP coupling weakens, while packing between HAMP subunits increases. Together, these changes loosen MH-cap structure, as shown by DEER measurements, reduce order in cryo-EM, and alter kinase control domain (KCD) dynamics. This change in flexibility propagates along the KCD helices to the CheA-CheW binding regions. Although isolated receptors may manifest these conformational shifts differently than the trimer-of-dimers organization in CheA–CheW complexes^12^, the upstream PAS/HAMP effects observed here are likely conserved as they are independent of higher-order array interactions.

Like the canonical Tar/Tsr receptors, Aer also possesses the TM, HAMP, and KCD regions that are directly involved in signal transmission. However, in Aer, conformational signals transmit from the flavin pocket through the PAS-MHL-cap interface to the KCD. In addition, Aer does not undergo adapative methylation^15^ despite retaining some conserved Glu or Gln residues in the analogous region. Sequence comparisons between Aer- and Tar-homologs are consistent with these differences (Fig. 3). For example, PAS interacting residues of Aer on the HAMP domain (L220, R244, Q248, L251) are generally not conserved by the Tar/Tsr receptors. Interestingly, MHL2 residues that interact with PAS in Aer show closer correspondence between Aer and Tar, with R497 highly conserved in both. In contrast, Aer does not conserve residues in MHL2 that are thought to be important for adaptation responses (e.g. the LLF motif and the poly-A track). In Tar/Tsr, the LLF motif stabilizes a zipped-helix cap structure that is critical for HAMP output control^30^. For Aer, the alternate VLL motif may facilitate PAS-MH interactions through increased flexibility. Loss of the VLL motif in Tsr abolishes signaling function of the receptor^27^, indicating that transduction to the KCD in Aer might differ from typical chemoreceptors as a result of direct KCD interactions with the sensory module. Residues within the bulges of the two phase shifts are similar for the AS-2-MHL2 junction, but quite different for the TM-AS-1 junction, with the bulge-stabilizing residue R235, and the membrane-embedded W206 not conserved in Tar (Fig. 3). This difference may reflect the role of the phase shift in transmitting conformational signals in Tar, whereas in Aer, the TM and HAMP may act more as a scaffold for the PAS-MHL-cap signaling axis.

Helix bulges and changes in local helix packing could amplify to larger scale movements or changes in dynamics at the receptor tip, as has been observed for soluble MCP-like proteins^61^. PAS-MH interactions combined with changes in HAMP stability could couple to MHL1 bulging, inducing packing rearrangements at the phase stutter. Many studies have indicated that conformational signals propagate through the KCD of chemoreceptors through changes in the dynamics and stabilities of these units^10, 28, 32, 51, 62–65^, although the detailed nature of the ensembles that these activity states represent is not fully resolved, especially in the context of the higher order receptor arrays. The observation that the MHL-bound and MHL-free forms of Aer coexist in the oxidized Q state when the receptors are free from additional interactions provided by their native environment, but that MHL-free is still favored by Aer ASQ, agrees with recent single-molecule FRET experiments that show how signals shift between populations of conformational states within broad ensembles^7, 65^. These experiments demonstrate increased conformational heterogeneity in the C-terminal helix of Tar on receiving a kinase-off signal, similar to that observed in the DEER experiments on Aer, despite the labeling positions being somewhat different in the two studies^65^. Aer and LBD-containing MCPs share similar architectures and conserved components that must converge on the same effectors, hence, they likely share elements of their signaling mechanisms. Nonetheless, their sequences have also adapted and diverged in specific regions to allow common components to assume different roles based on different mechanisms of signal input.

## Methods

### Protein purification of Aer solubilized in LMNG

BL21-DE3 *Escherichia coli* cells were co-transformed with a pET28a vector containing N-terminally Twin-strep-tagged *E. coli* Aer and a duet pAYC vector containing untagged *E. coli* CheA and CheW. Overnight pre-cultures were grown at 37 °C while shaking. 1.333 L of terrific broth (TB) media with 10% glycerol was inoculated with 20 mL of overnight culture. Cells were grown at 37 °C while shaking until an OD_600_ of 0.4 was reached. The temperature was reduced to 16 °C and 25 mg/mL IPTG was added to induce overexpression of Aer when the OD_600_ reached 0.6. Protein was expressed at 16 °C while shaking overnight.

Cells were harvested through centrifugation at 5000 RPM (Beckman Coulter JLA 9.100) for 10 min at 4 °C. Per 1.333 L of culture, a cell pellet of 10-15 mL was collected. 25-30 mL of cell pellet was thawed on ice and resuspended in lysis buffer (50 mM Tris pH 8.0, 200 mM KCl, 10% glycerol) to a final volume of 100 mL. The suspension was incubated with 60 mg lysozyme and 0.1 mM PMSF for an hour at 4 °C and sonicated on ice. Lysate was first centrifuged at 12,000 x g RPM (Beckman Coulter JA 30.5) for 45 min at 4 °C to remove cell debris and then ultracentrifuged at 100,000 x g for 1 h at 4 °C to acquire the insoluble fraction. The insoluble fraction was gently resuspended in lysis buffer at a 1:2 ratio. 1 % Lauryl Maltose Neopentyl Glycol (LMNG, Anatrace) was added to the resuspension diluted in equal volume lysis buffer and incubated overnight at 4 °C while rocking.

Ultracentrifugation at 100,000 x g for 1 h at 4 °C removed any remaining insoluble material. Affinity chromatography was performed using 5 mL Strep-Tactin®XT resin (IBA Lifesciences). Bound protein was washed 3 times with wash buffer (25 mM Tris, 150 mM NaCl, 10% glycerol, 0.1% LMNG, pH 8.0) and then eluted with elution buffer (100 mM Tris, 200 mM NaCL, 100 mM biotin, 0.1% LMNG) at 4 °C. The eluted protein was concentrated down to a volume of 10 mL and run on a Cytiva HiLoad^TM^ 26/60 Superdex 200^TM^ prep grade column in size exclusion chromatography (SEC) buffer (25 mM Tris pH 8.0, 150 mM NaCl, 5% glycerol, 0.002% LMNG) at 4 °C. Fractions containing solubilized, purified *E.coli* Aer bound to FAD were concentrated down to 2-3 mg/mL at 4 °C, flash frozen, and stored at -80 °C.

### Aer mutants for site-directed spin labelling

To produce mutants for MTSL site-directed spin labelling, the three native cysteine residues in Aer were replaced as follows (C193S, C203S, C253L) to produce Aer Δcys. Subsequently, single residues placed along the HAMP and KCD domains were mutated to cysteine to ensure site-directed labelling by MTSL (A223C, S271C, Q300C, S324C, S347C, E375C, S414C, A437C, S461C, S488C). Modelling of dimeric Aer and rationalization of structural and dynamical changes was accomplished with the use of AlphaFold2-Multimer. The mutation D68V was subsequently made in these constructs to lock the receptor in a constitutive kinase-inhibiting conformation. PDS ESR measurements were used to probe position 253 as well using Aer C193S C203S ± D68V. Aer and the Aer Δcys variants used for ESR characterization were expressed in BL21(DE3) chemically competent cells using a kanamycin-resistant pet28a vector. Aer expressed in UU2611, UU2612, and UU2632 chemically competent cells was used for mass spectrometry analysis. The cells were grown in Terrific Broth and protein expression induced using 1 mM IPTG at an OD_600_ of 0.6-0.8. Following 16-20 hrs of expression, the cells were pelleted by centrifugation and frozen. Cell pellets were solubilized and sonicated for 6 minutes in 50 mM Tris, 500 mM KCl, 10% glycerol supplemented with 1% DDM, 1 mM PMSF and 100 μg/mL FAD. All buffers used in the purification were chilled to 4 °C. The lysate was clarified by centrifugation at 12,096 xg for 1 hr at 4 °C. The supernatant was added to 5 mL of High-Density Cobalt Agarose Beads preequilibrated with extraction buffer and gently inverted at 4 °C overnight. The agarose beads were then filtered in a column and washed with 25 mM Tris, 150 mM NaCl, 10% glycerol, pH=8.00 supplemented with 100 μg/mL FAD and 0.1% DDM. An additional 1 mL of chilled buffer supplemented with 10 uL of freshly prepared 100 mM MTSL was added, the column sealed, wrapped in tinfoil, and gently rocked at 4 °C overnight. Aer WT and Aer D68V used for functional phosphotransfer assays and mass spectrometry analysis were not subjected to MTSL labelling, nor were they incubated an additional day. The protein was then eluted with 25 mM Tris, 150 mM NaCl, 300 mM imidazole, 10% glycerol, pH=8.00 supplemented with 100 μg/mL FAD and 0.1% DDM. An FAD standard curve was used to calculate the concentration of Aer by measuring flavin absorbance at 450 nm.

### CheA, CheW, CheY, and MSP1D1

For purification of *E. coli* CheA, CheW, CheY, and Membrane Scaffold Protein 1D1 (MSP1D1 to compose the nanodiscs), pET28a plasmid containing the target construct and an N-terminal poly-histidine tag (6xHis) was transformed into BL21-DE3 *E. coli* cells. Overnight precultures of 100 mL of LB broth were grown at 37 °C. 20 mL of the overnight culture was inoculated into 2 L of LB broth and grown at 37 °C until an OD_600_ of 0.4 was reached. The temperature was reduced 17 °C and monitored until an OD_600_ of 0.6 was reached. To induce over-expression of the proteins, 25 mg/mL of IPTG was added. Protein was expressed at 17 °C while shaking overnight.

The cells were harvested through centrifugation at 4,700 x g (Beckman Coulter JLA 9.1000) for 10 min at 4 °C. The cells were resuspended in ∼30 mL of chilled lysis buffer (50 mM Tris pH 7.5, 150 mM NaCl, 5% glycerol) and lysed using a probe sonicator on ice. MSP1D1 was solubilized in 1% Triton X-100 during sonication. The lysate was then centrifuged at 48,000 x g (Beckman Coulter JA 30.5) for 45 minutes at 4 °C to remove cellular debris. The supernatant was run through a gravity purification column containing ∼3 mL of Nickel-NTA resin (Thermo Fisher) at 4 °C. After washing with chilled wash buffer (50 mM Tris pH 7.5, 150 mM NaCl, 20 mM Imidazole, 5% glycerol), bound protein was eluted with elution buffer (50 mM Tris pH 7.5, 150 mM NaCl, 200 mM Imidazole, 5% glycerol). Eluted protein was concentrated for injection on a Cytiva HiLoad^TM^ 26/60 Superdex 200^TM^ prep grade column in size exclusion chromatography (SEC) buffer (25 mM Tris pH 7.5, 150 mM NaCl, 5% glycerol) at 4 °C. SEC fractions that contained purified protein were concentrated to ∼10 mg/mL using a 10 or 50 kDa MWCO Amicon Ultra-15 Centrifugal Filter. The resulting protein solutions were aliquoted, flash frozen, and stored at -80 °C.

### Nanodisc Reconstitution of Aer

Following purification of Aer WT or its variants as described above, the protein was concentrated to as high a degree as possible. To prepare Aer nanodiscs, stoichiometric ratios of 1:5:50 Aer:MSP1D1:DMPC were combined on ice to a final sample volume of 500 μL and a final glycerol concentration of 10% (v/v). First, a solution of 20 mM sodium cholate, DMPC lipids, and supplemental glycerol was prepared and incubated for 15 minutes. Aer was then added, and the sample incubated for another 15 minutes before addition of MSP1D1. This solution was placed on a sample rocker at 4 °C for 45 minutes, after which 400 μL of hydrated Bio-Beads were added. Incubation at 4 °C proceeded for another 60 minutes, at which time the liquid sample was extracted, filtered through a 20 μm filter, and injected onto a Superose 6 analytical column stored at 4 °C and preequilibrated with 25 mM Tris pH 8.0, 150 mM NaCl, 10% glycerol. Fractions containing Aer, as determined by 450 nm absorbance, were pooled and concentrated.

### Oxidized Aer cryo-EM sample preparation and collection

A 4 μl volume of 0.3 mg/ml oxidized Aer was applied to PELCO easiGlow™ (Ted Pella) plasma cleaned grids (Quantifoil R 1.2/1.3 Cu 200 Mesh). Grids were blotted for 3 s at 4 °C at 90% humidity after 30 s sample incubation and vitrified in liquid ethane using a Vitrobot Mark IV (FEI). Samples were imaged using SerialEM automated data acquisition on a Talos Arctica (ThermoFisher Scientific) operating at 200 kV, equipped with a Gatan K3 detector and Gatan energy filter set at 20 eV slit width. 4206 movies were acquired at a nominal magnification of 79,000x, corresponding to a pixel size of 1.04 Å/px. The total dose per exposure amounted to 50 e^-^/Å^2^, fractionated into 50 frames, with a defocus range from -0.6 µm to -2.2 µm.

### Oxidized Aer cryo-EM data processing

Single particle analysis was performed using cryoSPARC (v4.0 and later)^66^. Movies were imported for Patch Motion Correction and Patch CTF Estimation (Supplement). Initial blob picking and 2D classification allowed for Topaz training. Particles were picked by multiple rounds of Topaz training and extraction. Selected particles were used for Ab Initio Reconstruction and subsequent Heterogeneous Refinement, generating a particle set of 201,412 particles. Particle poses were refined using Non-Uniform Refinement^67^. This map was used as input for Reference-Based Motion Correction, resulting in 103,608 particles. Motion corrected particles were further refined through Non-Uniform Refinement and Local Refinement with imposed C2 symmetry and a refinement mask masking out the LMNG detergent ring. All Aer structures exhibited C2 symmetry even when refined without symmetry imposed. In parallel, a different particle set of 14,486 particles was selected in 2D based on the presence of defined KCD density, corresponding to an MHL-bound conformation of Aer. These particles were refined similarly with imposed C2 symmetry, reaching a global resolution of 4.0 Å. To separate the two populations, the MHL-bound particles were subtracted from the larger particle set list using the Particle Sets tool. The resulting 90,018 MHL-free particles were refined using Local Refinement with imposed C2 symmetry. The final map reached a global resolution of 3.5 Å.

### Reduced Aer cryo-EM sample preparation and data collection

Prior to vitrification, purified Aer sample was degassed in a mixed hydrogen (3%) and nitrogen (97%) atmosphere inside a Coy chamber and incubated with 1 mM dithionite. A 4 μl volume of 0.5 mg/ml reduced Aer was applied to PELCO easiGlow™ (Ted Pella) plasma cleaned grids (Quantifoil R 1.2/1.3 Cu 200 Mesh). Grids were blotted for 3 s at 6 °C at 90% humidity after 30 s sample incubation and vitrified in liquid ethane using a Vitrobot Mark IV (FEI) inside the Coy chamber. Data was collected at the Rutgers CryoEM & Nanoimaging Facility (RCNF) using automated data acquisition with EPU on a Titan Krios (Thermo Fisher Scientific) operating at 300 kV. A Gatan K3 direct electron detector was used in counting mode, equipped with a BioQuantum-LS energy filter set to a 20 eV slit width. A total of 11,975 movies were recorded at a nominal magnification of 130,000x. Each exposure was acquired with a total dose of 54.25 e⁻/Å², fractionated into 40 frames, at a nominal defocus range of -0.8 to - 2.4 μm.

### Reduced Aer cryo-EM data processing

11,975 movies were imported to cryoSPARC (v4.0 and later)^66^, followed by Patch Motion Correction and Patch CTF Estimation. Blob picking and 2D classification was used to generate an initial particle selection for Topaz training. Multiple rounds of Topaz training, extraction, and 2D classification resulted in a particle set of 202,990 particles. Particle poses were refined using Non-Uniform Refinement^67^. The resulting map was used as input for Reference-Based Motion Correction. Heterogeneous refinement of motion-corrected particles resulted in a final particle selection of 77,320, which refined to a global resolution of 3.3 Å using Non-Uniform Refinement and Local Refinement with imposed C2 symmetry and a refinement mask masking out the density of the LMNG detergent micelle.

### Structure refinement

Models of oxidized and reduced Aer were obtained by using an AlphaFold3 generated in silico model. Rigid body fitting in ChimeraX^68^ and subsequent molecular dynamics fitting using ISOLDE^69^ resulted in models suitable for real space refinement in PHENIX. Final models were acquired through iterating between manual model building in COOT^70^ and real space refinement against the cryo-EM density maps in PHENIX^71^. MOLPROBITY^72^ was used for validation of the structures.

### UV-Vis spectrometry

Purified Aer at ∼2 mg/mL was diluted to 0.3-1.5 mg/mL at a final volume of 100 µL in SEC buffer (25 mM Tris pH 8.0, 150 mM NaCl, 5% glycerol, 0.002% LMNG). Using a quartz cuvette with a 1 cm width, the UV-visible absorbance spectrum was measured from 300-800 nm at 10 °C. The sample was then incubated in an anaerobic glove chamber for 30 minutes at 4 °C to remove dissolved oxygen from the solution. The protein was reduced with the addition of dithionite (dissolved in 1M Tris pH 8.0) to a final concentration of 1 mM or 10 mM and incubated for 30 minutes at 4 °C. For some samples, removal of dithionite was accomplished by buffer exchange into SEC buffer in the anaerobic chamber through several rounds of dilution and concentration using a 0.5 ml centrifugal filter with a 50 kDa MWCO at 4 °C. The reduced protein sample was placed in a pre-cooled quartz cuvette and capped with a rubber septum while in the anaerobic chamber. The UV-visible absorbance spectrum of reduced Aer was measured from 300-800 nm at 10 °C. The septum was then removed, and Aer was re-oxidized with gentle stirring of the cuvette. The absorbance spectrum of re-oxidized Aer was measured and compared to the oxidized and reduced samples.

### FAD quantification and binding assays

Purified Aer at various concentrations (1.5-0.15 mg/ml) was prepared in SEC buffer. Protein concentrations were determined using Bradford reagent. Using a UV/Vis spectrophotometer set at 10 °C, the absorbance at A_445_ was measured for each sample, which corresponds to the absorbance of oxidized FAD. Using the A_445_ FAD extinction coefficient (11.3 mM^-1^ cm^-1^), the concentration of oxidized FAD in the protein sample was calculated and compared to the concentration of Aer subunit.

### Continuous wave electron paramagnetic resonance

Purified Aer (1.3 mg/ml) was incubated in an anaerobic chamber for 30 minutes at 4 °C. The protein was then reduced with the addition of 1 mM dithionite and incubated for 30 minutes at 4 °C. Dithionite removal was accomplished by buffer exchange into SEC buffer in the anaerobic chamber through several rounds of dilution and concentration using a 0.5 ml centrifugal filter with a 50 kDa molecular weight cuttoff at 4 °C. The removal of dithionite occurred over two hours. Samples were then prepared in the anaerobic chamber by pipetting 15 µl of the samples into pre-cooled EPR capillary tubes, which were then sealed using a two-component epoxy. CW-ESR spectra were collected at 10 °C on Bruker Elexsys E500 EPR instrument at 9.4 GHz with 100-kHz modulation frequency and 1.5- or 2-G modulation amplitude. After data collection, the protein concentration of each sample was determined using Bradford reagent and the resulting spectra were scaled appropriately.

### PDS-ESR Measurements

For 4-pulse-DEER, the samples were exchanged into deuterated buffer containing 30% d^8^-glycerol, checked by cwESR and plunge frozen into liq. N_2_. The pulse ESR measurements were carried out at Q-band (∼34 GHz) on a Bruker E580 spectrometer equipped with a 10 W solid state amplifier (150W equivalent TWTA) and an arbitrary waveform generator (AWG). DEER was carried out using four pulses (π/2-τ_1_-π-τ_1_-π_pump_-τ_2_-π-τ_2_-echo) with 16-step phase cycling at 60 K. The pump and probe pulses were separated by 56 MHz (∼20 G). The DEER data was background subtracted and the distance reconstruction was carried out using denoising and the SVD method^73^. Time-domain data for all PDS-ESR measurements can be found in Fig. S9.

### CheA autophosphorylation with LMNG-solubilized Aer

For the CheA autophosphrylation assays with Aer solubilized in LMNG, 25 µL samples containing 2 µM CheA and 2 µM CheW were incubated at 4 °C in the presence or absence of 6-12 µM Aer in TKEDM buffer (10 mM Tris pH 7.5, 100 mM KCl, 5 mM EDTA, 1 mM DTT, 5 mM MgCl_2_) supplemented with 0.002 % LMNG and 5 % glycerol. A mixture of 1 mM ATP mixed with radiolabeled γ-^32^P ATP at a final reactivity of 0.15 +/- 0.05 mCi was added to each reaction in a 2 µL volume. After 1 minute, the reaction was quenched with 25 µL of 3x LDS buffer containing 100 mM EDTA. 40 µL of each sample were ran on a native Tris-glycine gel at 120 volts for 2 hours using chilled native Tris-gly running buffer. The gels were dried overnight, placed in a radiocassette for a minimum of 24 hours, and then imaged with a Typhoon phosphor-imager. Image J software was used to quantify the relative intensities of the CheA-^32^P bands. For the CheY phosphor-transfer assays, 50 µM was added to the samples described above. The resulting CheY-^32^P band was quantified using Image J software. For reduced samples, dithionite dissolved in 1 M Tris pH 8 was added to each sample at a final concentration of 1 mM while in an anaerobic Coy chamber. The samples were incubated in the glove chamber at 4 °C for a minimum of 10 minutes before the assay was conducted.

### CheA autophosphorylation with Aer in nanodiscs

CheA autophosphorylation and phosphotransfer to CheY was monitored by ^32^P incorporation. All radioisotope assays were carried out in 50 mM MOPS, 150 mM KCl, 10 mM MgCl_2_, 10% glycerol, pH=7.5. Samples were prepared with 2 μM CheA (consistent within each experiment), 4 μM CheW, 50 μM CheY, and ∼6 μM Aer WT / Aer D68V in nanodiscs at a final volume of 23 μL. Following room temperature incubation for ∼15 minutes, [γ-^32^P]ATP was added to a final concentration of 1 mM and the reaction was quenched after 30 seconds using 4x SDS-PAGE loading buffer containing 50 mM EDTA, pH 8.0. The samples were loaded onto a 4-20% Tris-glycine polyacrylamide protein gel purchased from Invitrogen. Gel electrophoresis was carried out for 35 minutes at 125 V constant voltage using a running buffer supplemented with 10 mM EDTA. The resulting gels were dried in a Bio-Rad Gel Dryer overnight and placed in a radiocassette for >20 hrs prior to imaging on a Typhoon Image Scanner. Gel bands were quantified using ImageJ.

## Acknowledgments

These studies were supported by NIH grant R35GM122535. We thank the National Center for CryoEM Access and Training (NCCAT), the Cornell Center for Material Science (CCMR) and the Rutgers CryoEM & Nanoimaging Facility for access to cryo-EM data collection facilities. ESR experiments were conducted at ACERT which is supported by the National Institute of General Medical Sciences of the National Institutes of Health under Award Number 1R24GM146107. We thank Chloe Holod and Nozomi Ando (Cornell University) for assistance with EM grid freezing under anaerobic conditions. Competing interests: The authors declare that they have no competing interests.

Data and materials availability: The cryoEM data and structures are available at the Protein Data Bank with accession codes: 31CK, 31CJ, and 31JS. All other data needed to evaluate the conclusions in the paper are present in the paper or the Supplementary Materials.

