## Supplementary material for "Signaling mechanism of the transmembrane energy receptor Aer": SI Appendix, Fig. S1A

| Dataset | Oxidized Aer | | Reduced Aer |
|---|---|---|---|
| State | MHL-free homodimer | MHL-bound homodimer | Homodimer |
| **Data Collection and Processing** | | | |
| **Microscope** | Arctica | | Krios |
| **Voltage (keV)** | 200 | | 300 |
| **Detector** | Gatan K3 | | Gatan K3 |
| **Nominal magnification** | 79,000 | | 130,000 |
| **Data Acquisition Software** | Serial EM | | EPU |
| **Electron dose ($e^{-}/Å^{2}$)** | 50 | | 54 |
| **Pixel Size (Å) (binned )** | 1.04 | | 0.64 |
| **Defocus range (µm)** | 0.6-2.2 | | 0.8-2.4 |
| **Number of movies (#)** | 4206 | | 11975 |
| **Number of particles** | 90,018 | 14,486 | 77,320 |
| **Symmetry imposed** | C2 | C2 | C2 |
| **Resolution (Å)** | 3.5 | 4.0 | 3.3 |
| **FSC threshold** | 0.143 | 0.143 | 0.143 |
| **Refinement** | | | |
| **Initial model used** | AlphaFold3 | AlphaFold3 | AlphaFold3 |
| **Atoms** | 8069 (Hydrogens: 4051) | 10666 (Hydrogens: 5338) | 8072 (Hydrogens: 4054) |
| **Protein residues** | 496 | 664 | 496 |
| **Ligand (#)** | FAD (2) | FAD (2) | FAD (2) |
| **RMS Deviation** | | | |
| **Bond lengths (Å)** | 0.003 | 0.003 | 0.005 |
| **Bond angles (°)** | 0.627 | 0.515 | 0.966 |
| **MolProbity score** | 1.71 | 1.88 | 1.49 |
| **Clashscore** | 10.16 | 11.16 | 7.19 |
| **Poor rotamers (%)** | 0.24 | 0.00 | 0.48 |
| **Ramachandran Favored (%)** | 96.95 | 95.43 | 97.56 |
| **Ramachandran Allowed (%)** | 2.85 | 4.57 | 2.44 |
| **Ramachandran Disallowed (%)** | 0.20 | 0.00 | 0.00 |
| **Fit to map (CCmask)** | 0.71 | 0.75 | 0.77 |
| **EMDB (maps)** | EMD-58290 | EMD-58450 | EMD-58291 |
| **PDB (model)** | 31CK | 31JS | 31CJ |

**Table S1. Cryo-EM data collection, refinement, and validation statistics.**

**Table S2**

| Aer Structural Element | Labelled Cysteines | Lower Distance Peak Gaussian Fits: Peak Maxima (Å) | | | σ (Å) | | | <40Å Average Distance (Å) | | |
|---|---|---|---|---|---|---|---|---|---|---|
| | | Aer Δcys | Aer Δcys D68V | *D68V - WT* | Aer Δcys | Aer Δcys D68V | *D68V - WT* | Aer Δcys | Aer Δcys D68V | *D68V - WT* |
| HAMP AS-1 | A223C | 31.6 | 31.6 | 0.0 | 1.19 | 1.37 | -1.8 | 29.8 | 29.0 | 0.8 |
| HAMP AS-2 | 253C | 31.3 | 31.6 | -0.3 | 1.31 | 1.32 | -0.1 | 28.9 | 29.2 | -0.3 |
| N-Terminal Helix | S271C | 31.9 | 31.6 | 0.3 | 1.37 | 1.29 | 0.8 | 29.3 | 29.6 | -0.3 |
| | Q300C | 29.7 | 29.7 | 0.0 | 1.21 | 1.19 | 0.2 | 28.5 | 28.4 | 0.1 |
| | S324C | 31.6 | 30.0 | 1.6 | 1.22 | 1.21 | 0.2 | 29.2 | 28.5 | 0.7 |
| | S347C | 34.3 | 31.6 | 2.7 | 1.58 | 1.31 | 2.8 | 29.5 | 29.4 | 0.2 |
| | E375C | 31.6 | 34.3 | -2.7 | 1.14 | 1.12 | 0.2 | 28.9 | 30.8 | -1.9 |
| C-Terminal Helix | S414C | 31.6 | 39.8 | -8.2 | 1.14 | 1.34 | -2.0 | 29.9 | 29.4 | 0.5 |
| | A437C | 30.2 | 31.6 | -1.4 | 1.28 | 1.35 | -0.7 | 28.8 | 29.3 | -0.5 |
| | S461C | 30.5 | 34.3 | -3.8 | 1.12 | 0.72 | 3.9 | 28.0 | 30.9 | -2.9 |
| | S488C | 27.2 | 42.4 | -15.2 | 1.06 | 1.79 | -7.3 | 27.3 | 31.4 | -4.1 |

| Peak Maxima shift to: | σ shift to: | Average Distance shift to: |
|---|---|---|
| Higher Distance | More Dynamic | Higher Distance |
| No Change | No Change | No Change |
| Lower Distance | Less Dynamic | Lower Distance |

**Table S2**: **DEER data summary.** P(r) peak Shifts refer to the difference in maxima of the lower-distance peaks from each P(r) distribution (D68V - WT). σ values refer to Gaussian fits of individual peaks. States with greater inter-subunit separations at a given position in Aer Δcys D68V sample relative to Aer Δcys sample are designated in red boxes. Shorter inter-subunit separations are specified with a cyan box. Similarly, greater dynamics at a given position in Aer Δcys D68V sample relative to Aer Δcys is inferred by the width of the distance distribution and shown in yellow, where states with lower inferred dynamics in Aer Δcys D68V are indicated with a gray box.

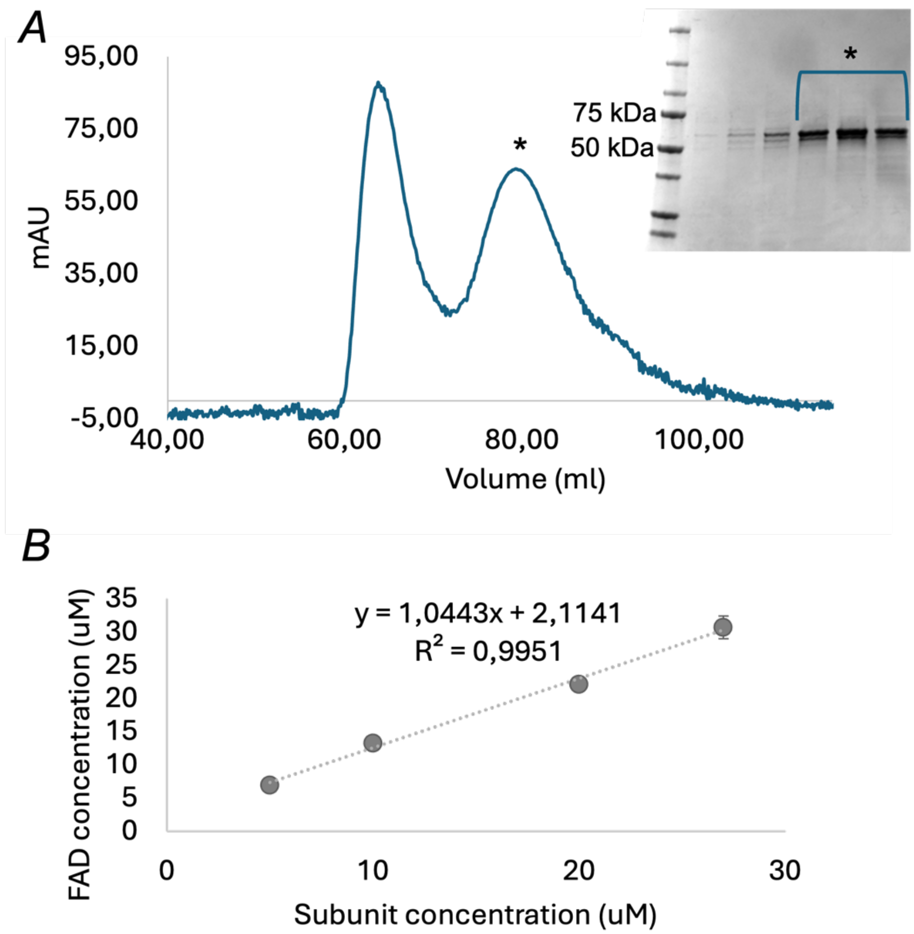


**Figure S1: Purification of Aer solubilized in detergent.** (A) Aer solubilized in the detergent LMNG was purified via affinity and size-exclusion chromatography (SEC). The SEC trace demonstrates that Aer elutes at a single peak. Inset shows an SDS-PAGE gel of the indicated elution peak stained with Coomassie dye. (B) UV/Vis spectroscopy analyses show that FAD is fully incorporated in detergent-solubilized Aer. The concentration of FAD was determined using the $A_{445}$ FAD extinction coefficient (11.3 $mM^{-1}$ $cm^{-1}$). The protein subunit concentration was determined using Bradford reagent.

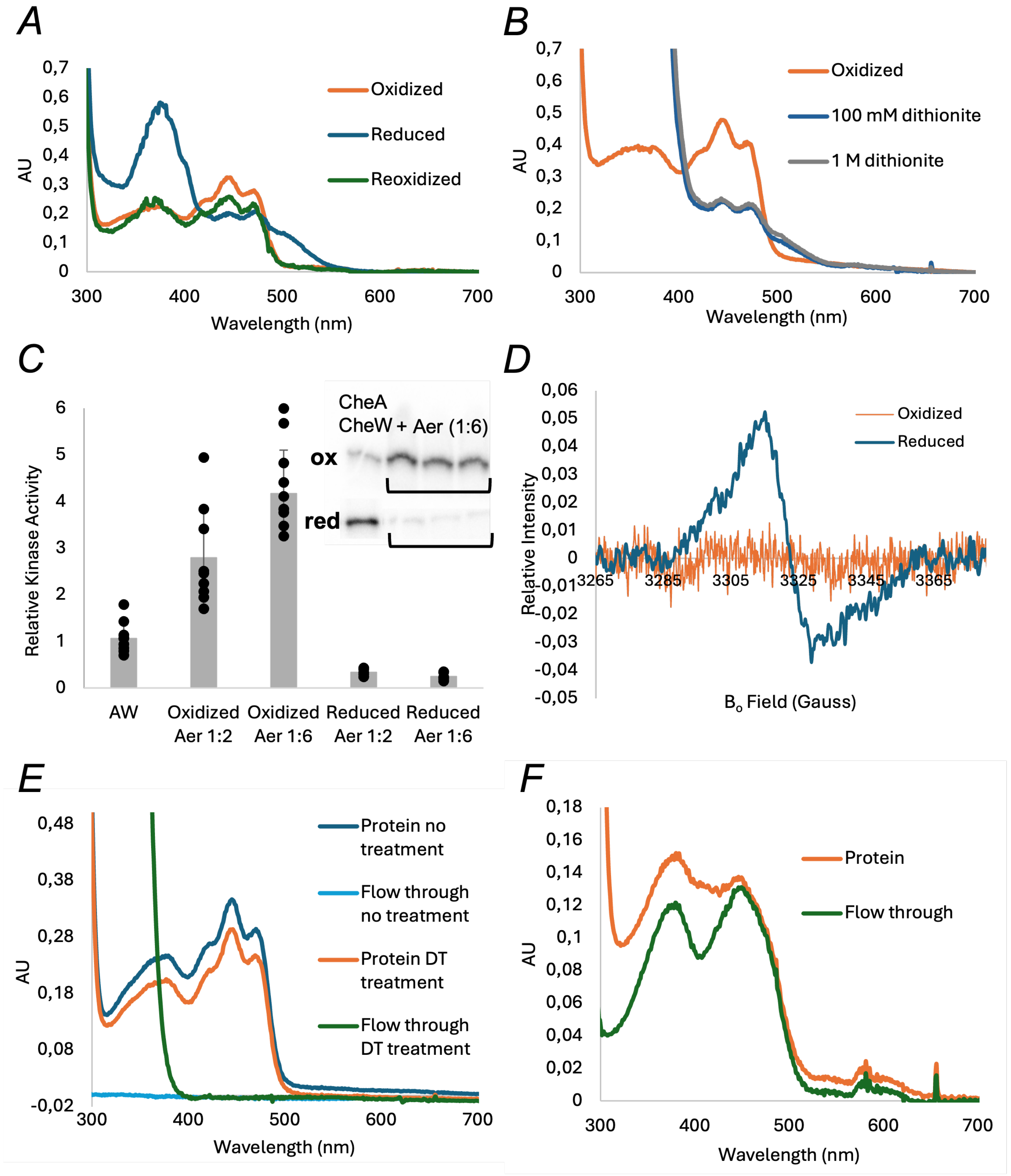


**Figure S2: Spectroscopic and redox properties of Aer.** (A) The redox state of FAD as monitored by UV/Vis spectroscopy. Aer that has been purified aerobically is bound to oxidized FAD (orange) that has signature absorption peaks at 365 nm, 416 nm, 441 nm, and 465 nm. When oxidized Aer is treated with dithionite under anaerobic conditions, there is a concurrent increase in absorbance at 372 nm and 500 nm, and a decrease of the peak at 441 nm with a slight shoulder at 395 nm, as indicative of the FAD anionic semiquinone (ASQ) state (blue). When the reduced ASQ is introduced to an aerobic environment, the cofactor is fully oxidized (green). (B) The addition of excess dithionite (100 mM – 1 M) does not reduce the FAD to the fully reduced hydroquinone (HQ) state. Instead, the ASQ state remains stable, which is evident by sustained absorbance peaks at 441 nm, 465 nm, and 500 nm. (C) Radioisotope assays that monitor CheA autophosphorylation show that oxidized Aer solubilized in LMNG detergent increases CheA autokinase activity. When Aer is present in a two-fold excess to CheA (1:2 ratio) there is a 2.8-fold increase in CheA activity. A six-fold excess of Aer (1:6 ratio) results in a 4.2-fold increase in CheA activity. Reduced Aer decreases CheA activity by 10-fold. Inset shows representative phosphorylation bands for oxidized (ox) and reduced (red) conditions. (D) Continuous wave electron spin resonance (ESR) experiments indicate the presence of an unpaired electron after the reduction of Aer with dithionite. (E) Reduction of FAD does not result in dissociation of the cofactor from the Aer subunit. Excess dithionite treatment under anaerobic conditions followed by centrifugal filtration through a 50 kDA MWCO filter demonstrates that FAD remains bound to Aer and is not present in the collected flow-through. (F) Treatment of Aer with 4 M guanidinium hydrochloride followed by centrifugal filtration through a 50 kDA MWCO filter demonstrates the dissociation of FAD from Aer.

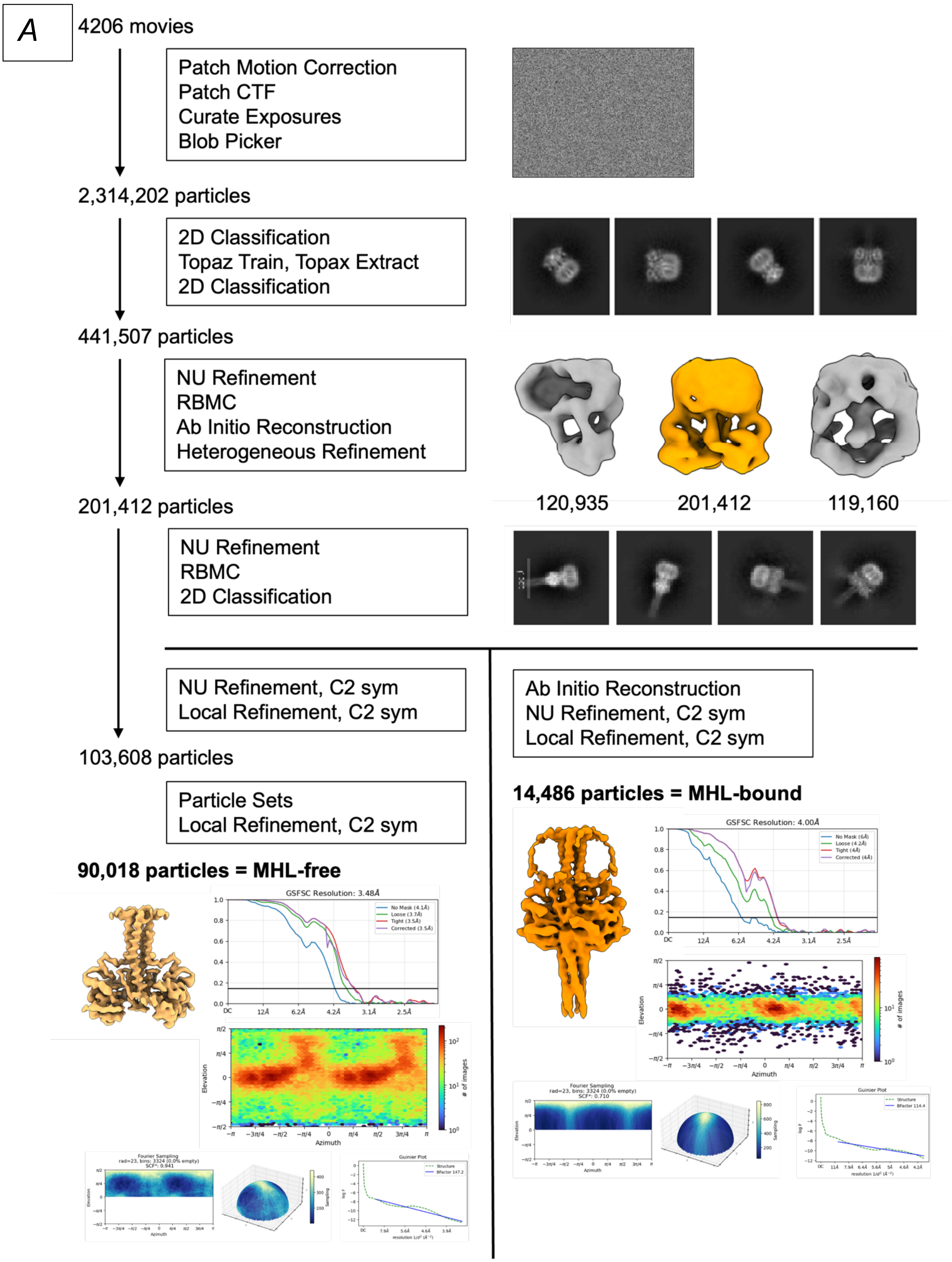
A
4206 movies
Patch Motion Correction
Patch CTF
Curate Exposures
Blob Picker
2,314,202 particles
2D Classification
Topaz Train, Topax Extract
2D Classification
441,507 particles
NU Refinement
RBMC
Ab Initio Reconstruction
Heterogeneous Refinement
120,935
201,412
119,160
201,412 particles
NU Refinement
RBMC
2D Classification
NU Refinement, C2 sym
Local Refinement, C2 sym
103,608 particles
Particle Sets
Local Refinement, C2 sym
90,018 particles = MHL-free
GSFSC Resolution: 3.48Å
Ab Initio Reconstruction
NU Refinement, C2 sym
Local Refinement, C2 sym
14,486 particles = MHL-bound
GSFSC Resolution: 4.00Å
Fourier Sampling
Guinier Plot

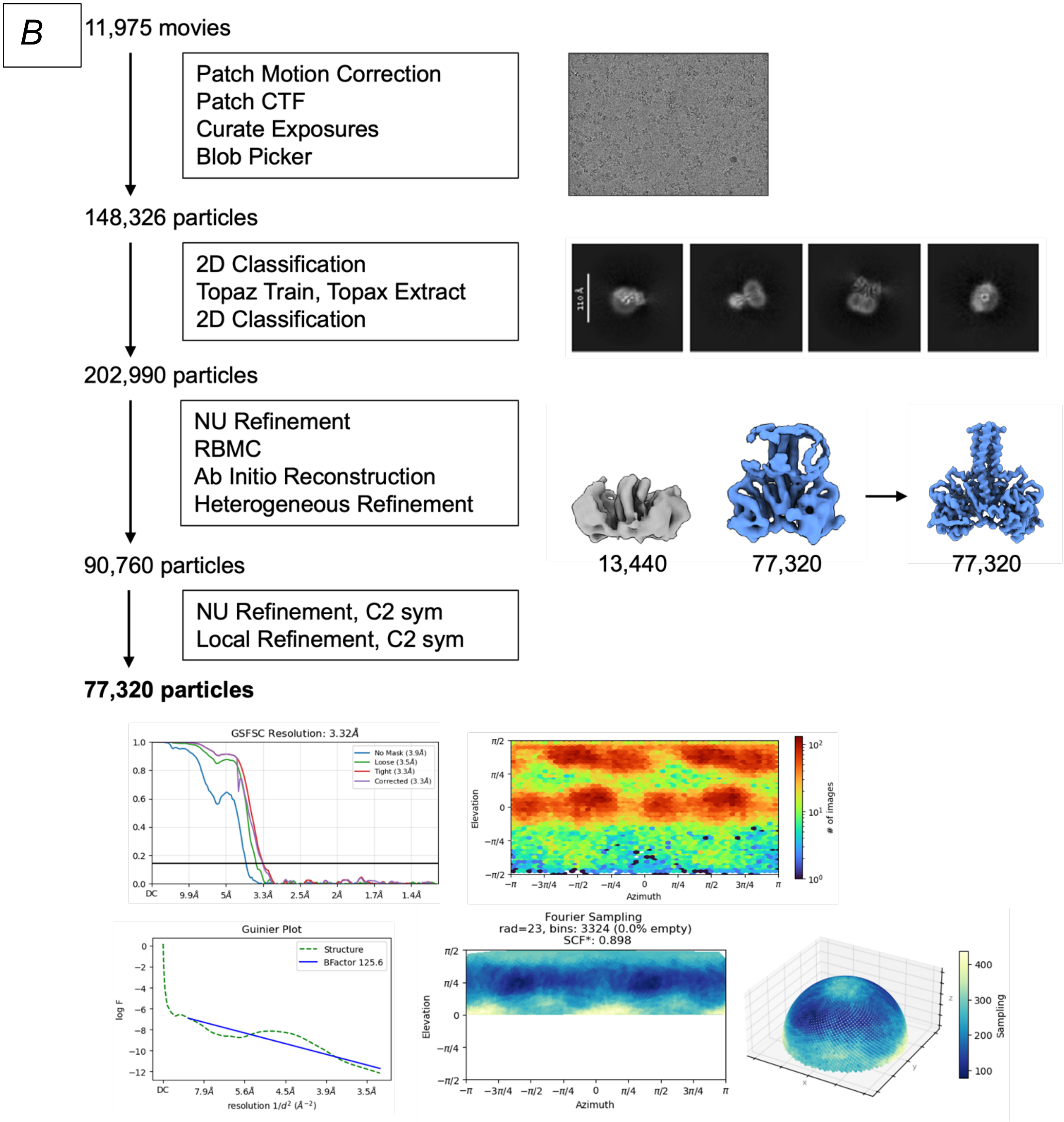


**Figure S3: Cryo-EM processing workflow for Aer in the MHL-free and MHL-bound Q state (A) and in the ASQ state (B).** The workflow, representative 2D and 3D classes as well as Gold-Standard Fourier Shell Correlation (GSFSC) curves, particle distribution, Guinier plots, and Fourier sampling of orientations are shown for both datasets. Particle numbers are shown for each processing sequence. CTF: contrast transfer function, NU: Non-Uniform Refinement, RBMC: Reference-Based Motion Correction.

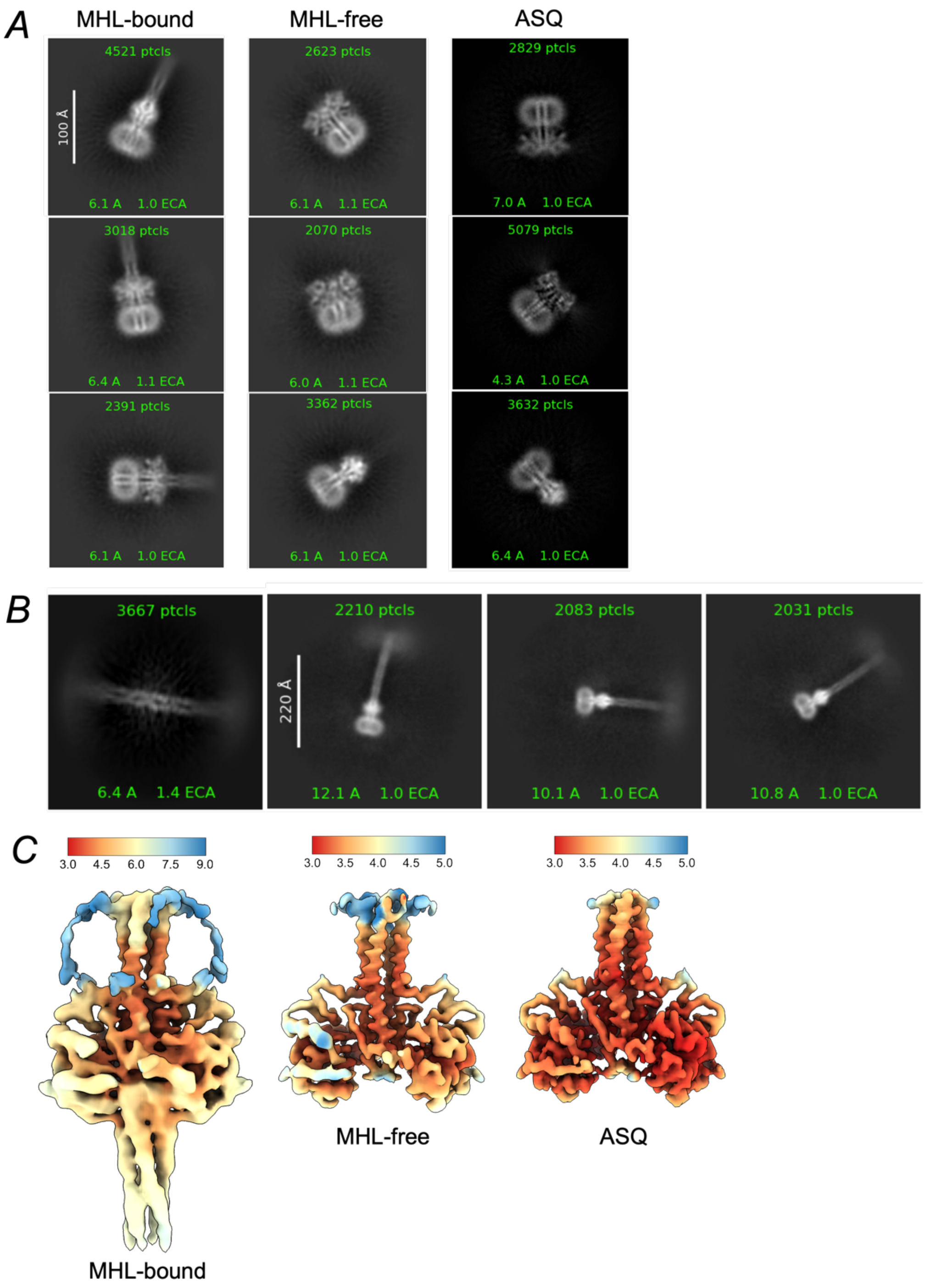


**Figure S4: Cryo-EM data of the different redox states of Aer.** (A) Representative 2D classes of Aer cryo-EM images in MHL-bound Q, MHL-free Q, and ASQ configurations. (B) MHL-bound 2D classes that resolve the KCD are only seen in the Q data set. (C) Local resolution data mapped onto Aer in MHL-bound Q, MHL-free Q, and ASQ states. The MHL-bound conformation is significantly lower resolution than the MHL-free and ASQ densities.

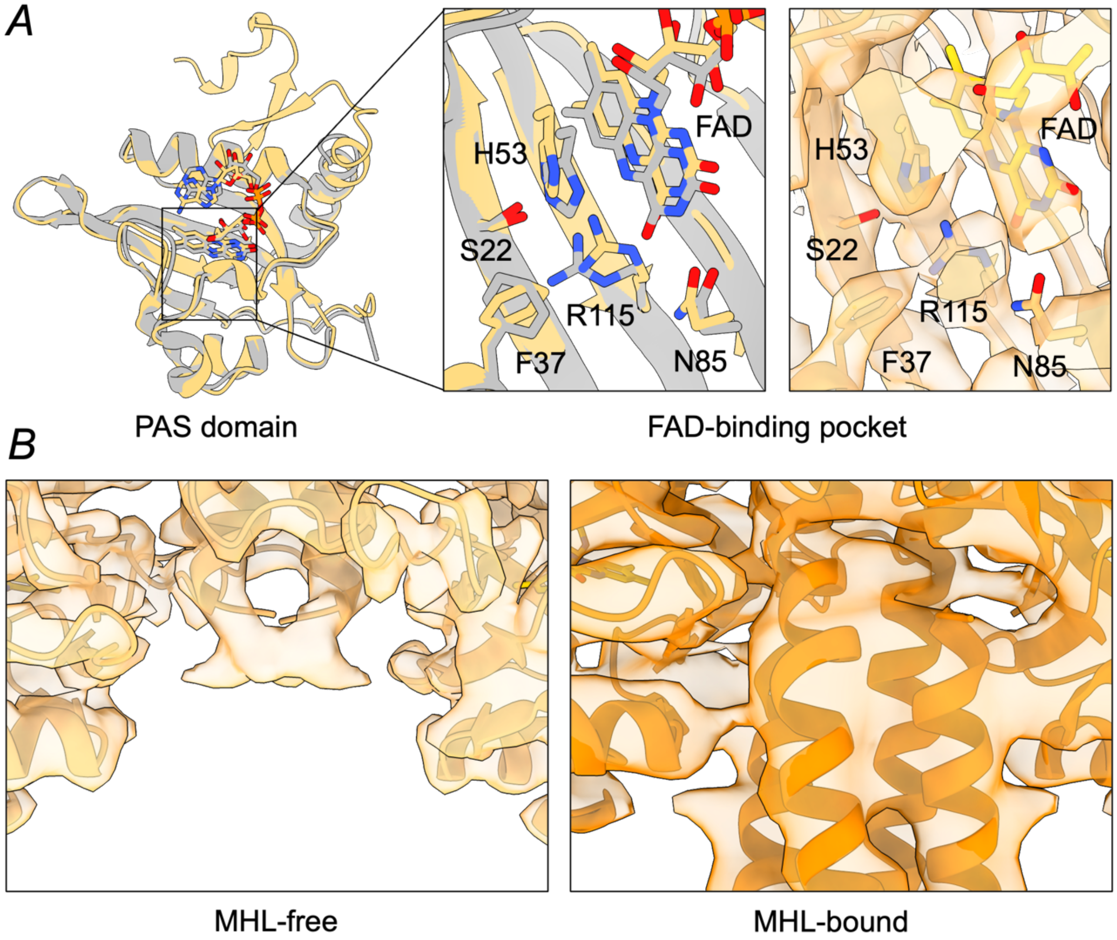


**Figure S5: Comparison of the Aer MHL-free structure with the isolated PAS-GVV structure.** (A) superposition of Aer MHL-free cryo-EM structure (tan) to Aer PAS-GVV crystallographic structure (gray, PDB ID 8DIK) in cartoon representation with expanded view of the FAD-binding pocket. Electron density for the FAD-binding pocket in Aer Q MHL-free is shown on the right. (B) Comparison of electron density in the MHL-free and MHL-bound Aer Q structures in the PAS-MHL interaction region.

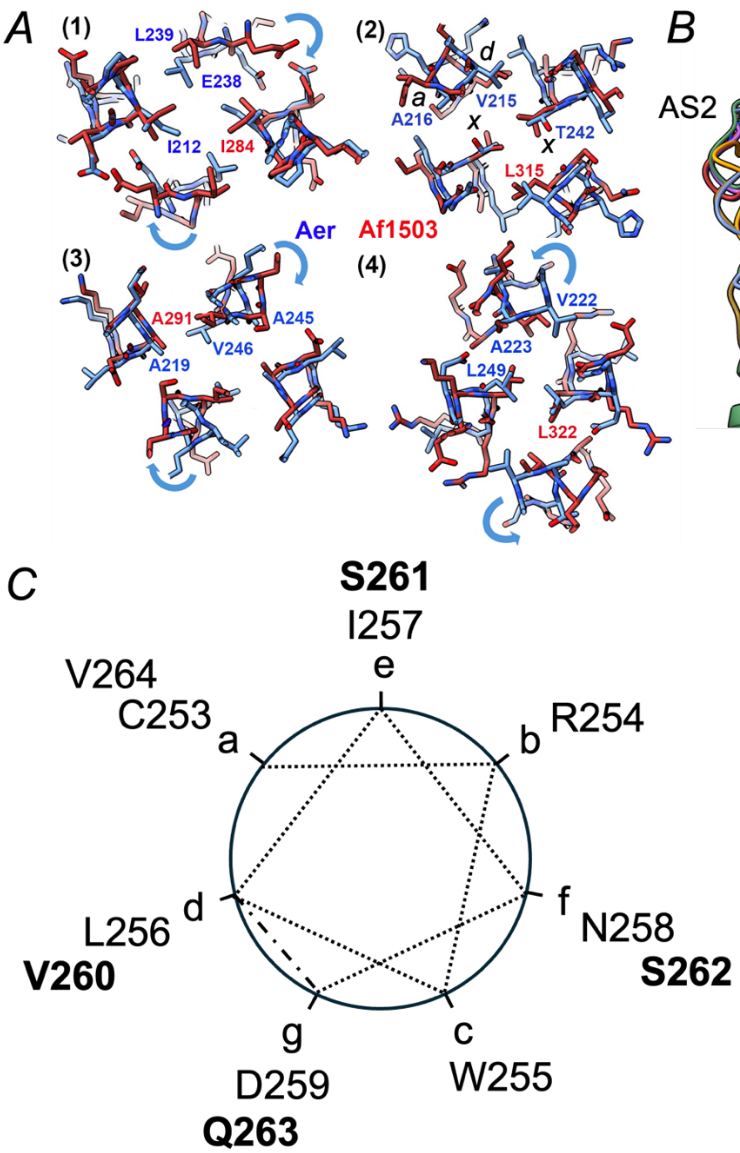


**Figure S6: Comparisons of the Aer HAMP conformation to those of other HAMP and HEAT motifs.** (A) Superposition of the four hydrophobic packing layers in the parallel bundle HAMP domain of Aer (blue) to that of Af1503 (PDB:2I7h, red), which displays x-da packing. Only layer 3 of Aer containing T242 as an "x" residue has close to x-da packing, with the other layers, more similar to knobs-into-holes packing of central "d" residues. Blue arrows depict how the opposed helices in Aer would have to rotate to assume x-da packing. However, there are additional differences in helix placements between the structures that make direct comparisons different. (B) Structural alignment between AS1 and AS2 of the Aer HAMP with other known HAMP structures reveal that in Aer AS2 (dark yellow) is displaced vertically half a helical step compared to the other domains (PDB ID: 2I7H, 4CTI, 3LNR, 5JEF, 5JEQ, 2Y0Q). Aer packing is more reminiscent of that observed in some 3-helix HEAT-repeat domains (8yab-B, gray) as revealed by a DALI search [56]. (C) Helical wheel schematic of the phase shift in the HAMP-MHL region using established heptad repeat nomenclature. Dotted lines represent typical helical packing, the dash-dotted line corresponds to the phase stammer at D259-V260.

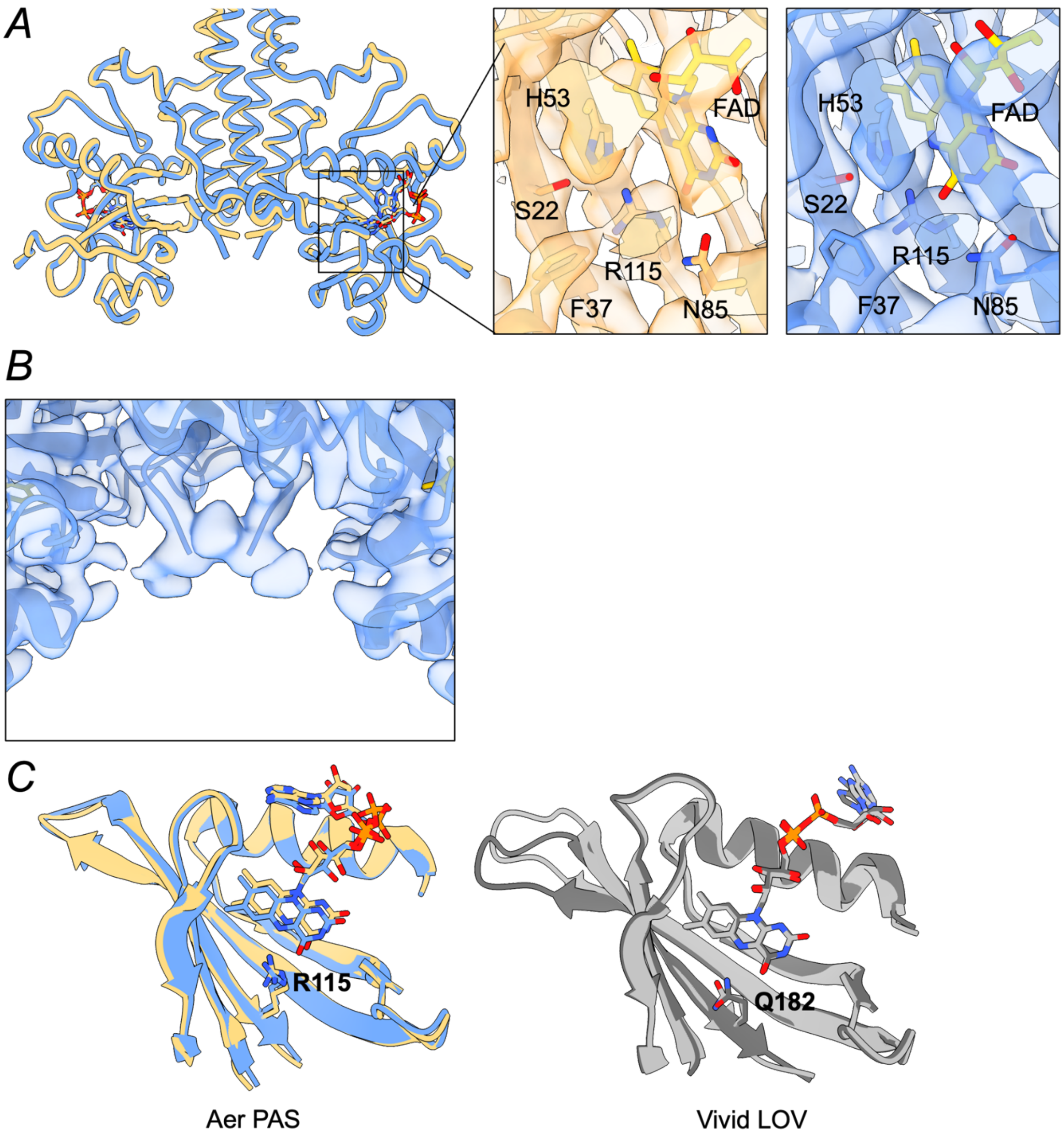


**Figure S7: Conformational changes upon Aer reduction to the ASQ.** (A) Overlay of the Aer MHL-free Q (tan) and Aer ASQ (blue) structures shown in cartoon representation. Inset shows electron density in the respective FAD-binding pockets. (B) Lack of electron density in the MHL-cap region of Aer ASQ. (C) Comparison between Aer PAS and the Vivid LOV domains in oxidized/reduced and light/dark state respectively. PDB ID: 3RH8 (light state, light gray), 2PD7 (dark state, dark gray).

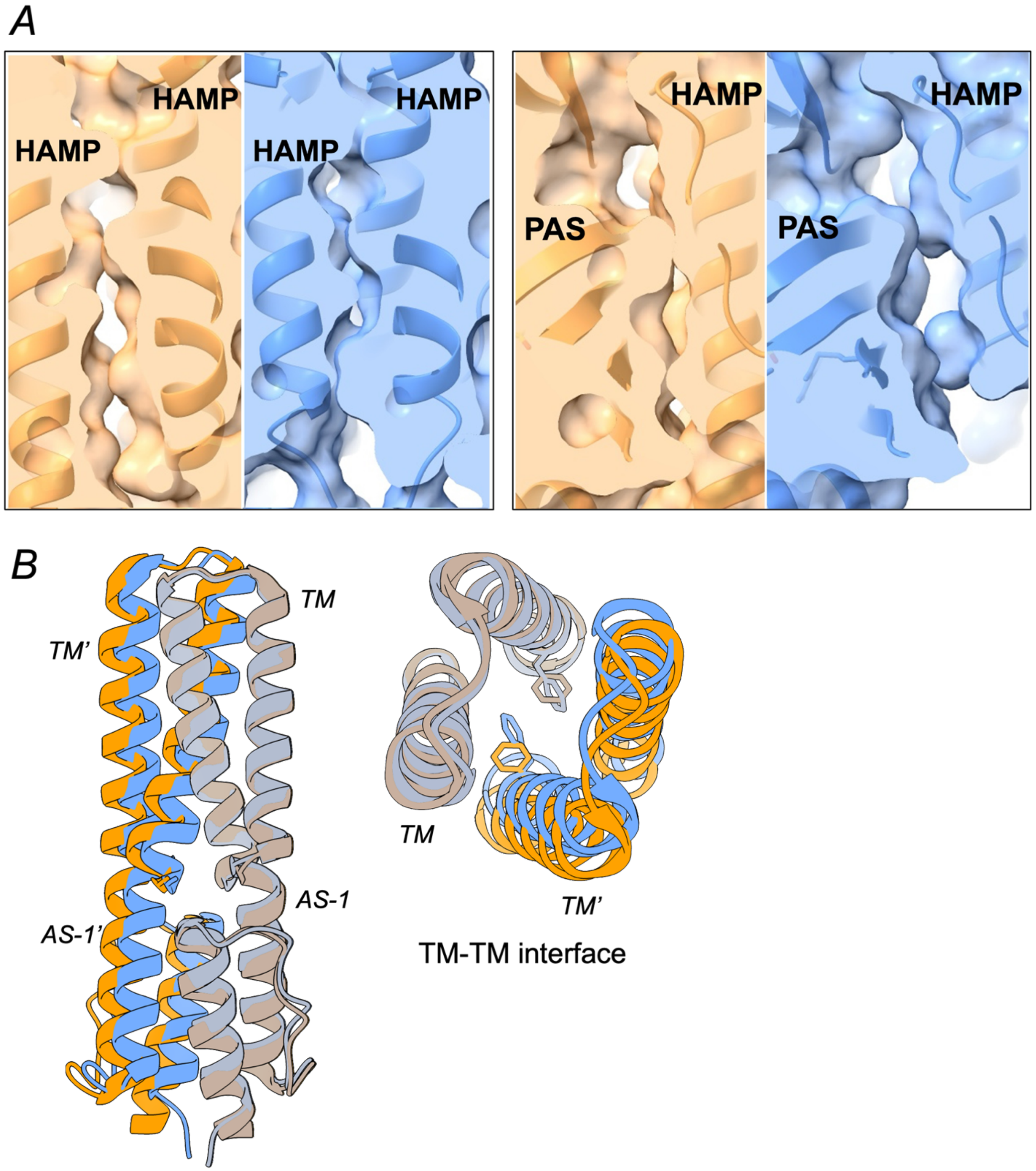


**Figure S8: Conformational changes upon Aer reduction to the ASQ.** (A) Packing changes at the HAMP-HAMP and PAS-HAMP interfaces in Aer MHL-bound Q (light orange) compared to Aer ASQ (blue) shown with both cartoon and surface representation. HAMP-HAMP contacts are looser and PAS-HAMP contacts tighter in the oxidized (Q) versus reduced state (ASQ) structures. (B) Changes in the TM-TM and HAMP-HAMP interface upon reduction from MHL-bound Q (orange) to the ASQ state (blue).

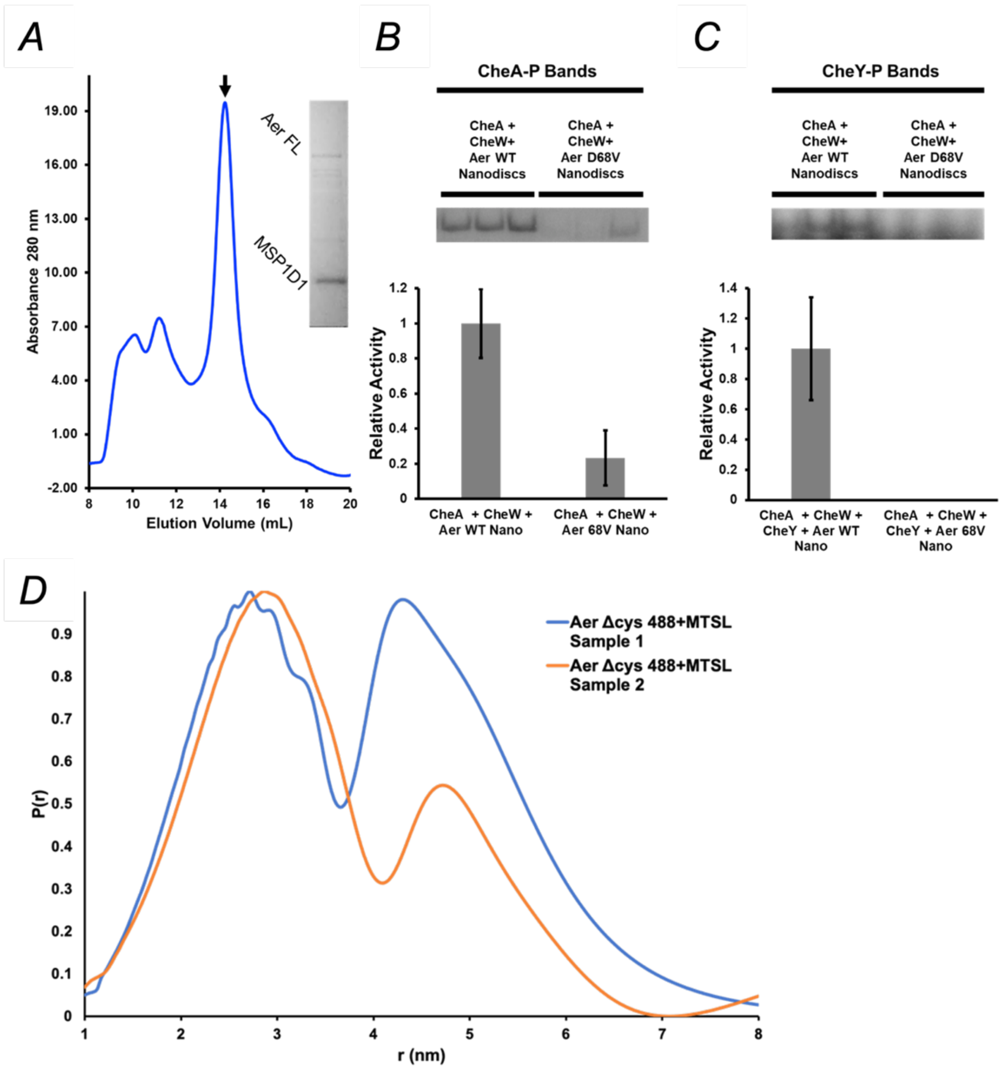


**Figure S9: CheA autophosphorylation and CheY phosphotransfer activity for nanodisc incorporated Aer.** (A) Aer incorporation into MSP1D1 Nanodiscs. Size-exclusion chromatogram of Aer in nanodiscs on an analytical S200 column. The inset Coomassie-stain SDS-PAGE of the indicated peak demonstrates Aer and MSP1D1 coelute. (B) Autophosphorylation of CheA in the presence of CheW, Aer WT nanodiscs, and Aer D68V nanodiscs. CheA-P (phosphorylated CheA) bands are shown above the quantification. CheA-P bands are normalized to the Aer WT in nanodisc condition. All experiments performed in triplicate; error bars are standard deviations. (C) Phosphotransfer to CheY in samples containing Aer WT and Aer D68V. CheY-P (phosphorylated CheY) bands are shown above the quantification. CheY-P bands are normalized to Aer WT in nanodisc condition. CheY-P bands from Aer D68V in nanodiscs are not detectable. All experiments performed in triplicate; error bars are standard deviations. (D) Reproducibility in P(r) distributions between separate sample preparations. Aer Δcys 488+MTSL was prepared twice, from separate growths in BL21/DE3 cells. Following purification, nanodisc incorporation, four-pulse DEER, background subtraction, denoising, and reconstruction using the SVD method, the resulting lower distance peaks were fit to Gaussian functions. The lower-distance peak maxima differ by 1.4 Å and their σ values differ by 0.36 Å. The longer-distance peaks differ by 4.0 Å and are at least partly variable owing to differences in disulfide-linked dimers.

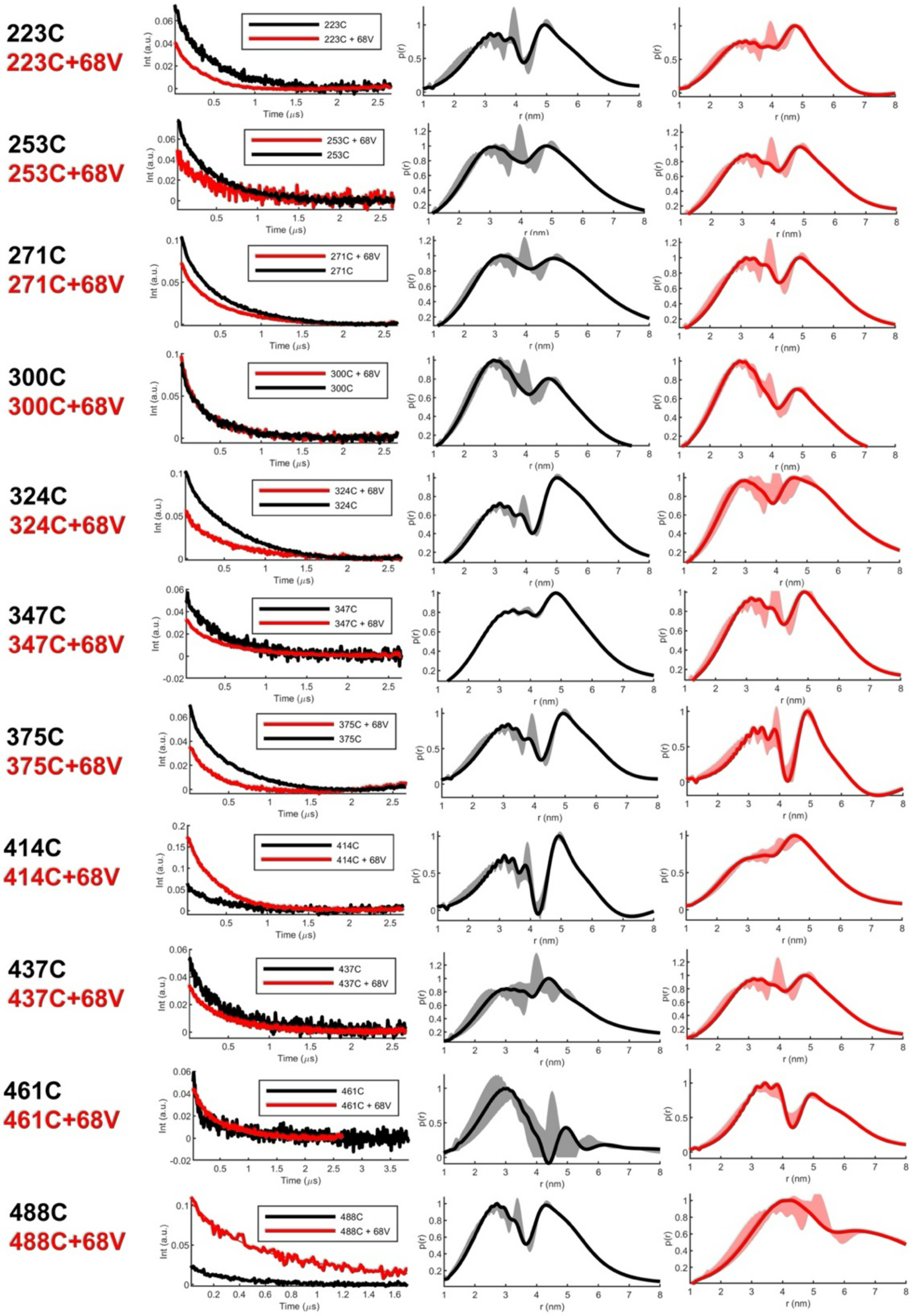


**Figure S10: DEER data for spin-labeled Aer.** Time-domain (left) and side-by-side distance distributions for 4P-DEER data on spin-labeled Aer at the positions indicated. Aer ∆cys shown in grey; Aer Dcys ∆68V in red. The distance domain was obtained by using the SF-SVD method. Errors in the distance distributions are represented by gray and red shading and calculated as described in [43].
